# Three transcription factors coordinate the archaeal cell cycle progression through a regulatory braking point mechanism

**DOI:** 10.1101/2025.10.16.681758

**Authors:** Yunfeng Yang, Shikuan Liang, Zixin Geng, Miguel V. Gomez-Raya-Vilanova, Wenying Xia, Junfeng Liu, Qihong Huang, Jinfeng Ni, Qunxin She, Mart Krupovic, Yulong Shen

## Abstract

Archaea of the order Sulfolobales execute a well-structured cell cycle program similar to that of eukaryotic cells. However, unlike eukaryotes, archaea lack cyclins and cyclin-dependent kinases and thus the mechanism of cell cycle regulation remained enigmatic. Here, we show that three essential ribbon-helix-helix domain transcription factors, namely, aCcr1, aCcr2, and aCcr3, play pivotal roles in controlling the cell cycle progression in the thermoacidophilic archaeon *Saccharolobus islandicus*. Coordinated expression of aCcr1 during M-G1 phase, aCcr3 during G1-S phase, and aCcr2 throughout the cell cycle ensures the timely transcription of the key genes that define the cell cycle phases. These include genes involved in chromosome dimer resolution, chromosome segregation, cell division, chromatin organization, DNA replication and repair, protein phosphorylation and degradation and metabolisms of amino acids, tRNA and carbohydrates. The synergy between aCcr1, aCcr2, and aCcr3 is achieved through their differencial affinities for the promoters and the levels of protein expression. We propose that the global regulation of the Sulfolobales cell cycle may be achieved not through transcriptional activation, but rather by repression of the key genes during strategic moments of the cell cycle. We propose a braking point model for the cell cycle control in Sulfolobales, which may represent a simple evolutionary intermediate on the way to the more complex cell cycle regulation in eukaryotes.

## Introduction

The cell cycle, the precisely coordinated sequence of events that governs the cell growth, DNA replication, and cell division, is essential for the proliferation of cellular organisms. Thermoacidophilic archaea of the order Sulfolobales encode many eukaryotic-like cell biology systems ^1,2^ and exhibit a eukaryotic-like cell cycle which is divided into well-defined phases ^1,3-7^. The cell cycle of an exponentially growing Sulfolobales cell starts with a short (∼5% of the cell cycle) pre-replicative phase, known as the G1 phase, during which the cell size increases in preparation for chromosome replication. The G1 is followed by the S phase (∼30-35% of the cell cycle), during which the genome is replicated. The G2 phase (∼50-60% of the cell cycle) is the second period of active cell growth, which is followed by the M phase, during which cells with two copies of aligned chromosomes undergo segregation, leading to cell division during the D phase. The M and D phases in Sulfolobales cells are short, each accounting for ∼5% of the cell cycle ^1^.

In eukaryotes, the cell cycle is regulated through balanced expression of cyclins and cyclin-dependent kinases (Cdks), with the expression of Cdks being relatively stable throughout the cell cycle and cyclins being synthesized and degraded in a cell cycle phase-dependent manner ^8,9^. The regulatory network of various cyclins and Cdks ensures that the cell cycle dependant waves of gene expression pass through numerous checkpoints. The three most critical checkpoints in eukaryotes are the G1 checkpoint, the G2/M checkpoint, and the spindle (M) checkpoint ^10,11^. In Sulfolobales, the processes of cell division, genome replication, repair, cell growth, and chromosome segregation have been the subject of considerable interest and are now relatively well understood ^2,12-20^. Recently, transcriptomic analysis of the cell cycle-synchronized *Saccharolobus islandicus* REY15A populations has shown that in addition to the core systems mentioned above, many other biological processes, including diverse metabolic pathways, protein synthesis, cell motility and even antiviral defense systems, are coordinated with the cell cycle ^21^. However, the regulatory mechanisms underlying the orderly transitions between different cell cycle phases remain largely unexplored. It is also unclear whether the archaeal cell cycle exhibits checkpoints similar to those observed during the eukaryotic cell cycle ^22^.

Recently, it has been reported that the proteasome controls the cell cycle progression by degrading CdvB, the major component of the cytokinetic ring in *Sulfolobus acidocaldarius* ^14^ and *Sa. islandicus* REY15A ^16^. This proteasome-dependent cell division regulation mirrors the mechanism described in eukaryotic cells, suggesting the existence of certain parallels between the cell cycle progression in Sulfolobales and eukaryotes ^14^. However, similar to other archaea, Sulfolobales lack identifiable genes for cyclins, suggesting differences in the cell cycle regulation compared to the cyclin/Cdk-dependent control operating in eukaryotes. Nevertheless, the transition between different cell cycle phases appears to be, at least partly, controlled at the transcriptional level. We and others have recently identified a ribbon-helix-helix (RHH) domain-containing transcription factor, aCcr1, which controls cytokinesis by repressing the expression of cell division genes ^23,24^.

Here, we describe two additional aCcr1 homologs encoded by *Sa. islandicus* REY15A. We show that the three aCcr transcription factors are essential to orchestrate a number of biological processes during the cell cycle progression. Specifically, aCcr1 and aCcr3 function during the M-G1 and G1-S transitions, respectively, whereas aCcr2 operates throughout the cell cycle. Based on our findings, we propose that cell cycle regulation in Sulfolobales occurs through a “braking point” mechanism, a model which resembles the eukaryotic checkpoints. We hypothesize that in nascent Sulfolobales cells, the promoters of genes important for cell cycle progression are innately active and are primarily regulated by the aCcr-mediated repression. De-repression of these genes at specific time points determines the smooth progression of the cell cycle. Our study forms a solid foundation for future exploration of the intricacies of cell cycle regulation in Sulfolobales and beyond.

## Results

### aCcr3 shows a cyclic expression, similar to that of aCcr1

We and others have recently reported that the cell cycle transcription regulator aCcr1 is expressed in *Sa. islandicus* exclusively during cytokinesis ^23,24^. Through transcriptomic and ChIP-seq analyses, we have identified the aCcr1-regulated target genes, including the cytokinesis initiation gene *cdvA.* It has been further predicted that *Sa. islandicus* species encode multiple aCcr1 homologs (Figure S1; 5 in *Sa. islandicus* REY15A and 4 in *Sa. islandicus* LAL14/1, including aCcr1) ^23^. This observation suggested that at least some of the aCcr1 homologs could be also involved in cell cycle regulation. Indeed, overexpression of SiL_RS14755 (hereafter referred to as aCcr2) in *Sa. islandicus* LAL14/1 led to growth retardation and cell enlargement ^23^. However, the exact functions and mechanism of action of aCcr2 (SiRe_1157) and other RHH domain-containing proteins, namely, SiRe_1577 (hereafter aCcr3), SiRe_1373, and SiRe_0131, in *Sa. islandicus* remained unknown.

RT-qPCR analysis showed that in exponentially growing (OD_600_=0.2) unsynchronized cells, the expression levels of aCcr1, aCcr2, and SiRe_1373 were rather similar, whereas the expression level of aCcr3 was approximately double that of the other homologs. By contrast, the expression level of SiRe_0131 was significantly lower, approximately one-tenth of that of aCcr1 (Figure 1A). These results are consistent with the transcriptomic analysis of synchronized cells ^24^ (Figure S2A).

**Figure 1.**
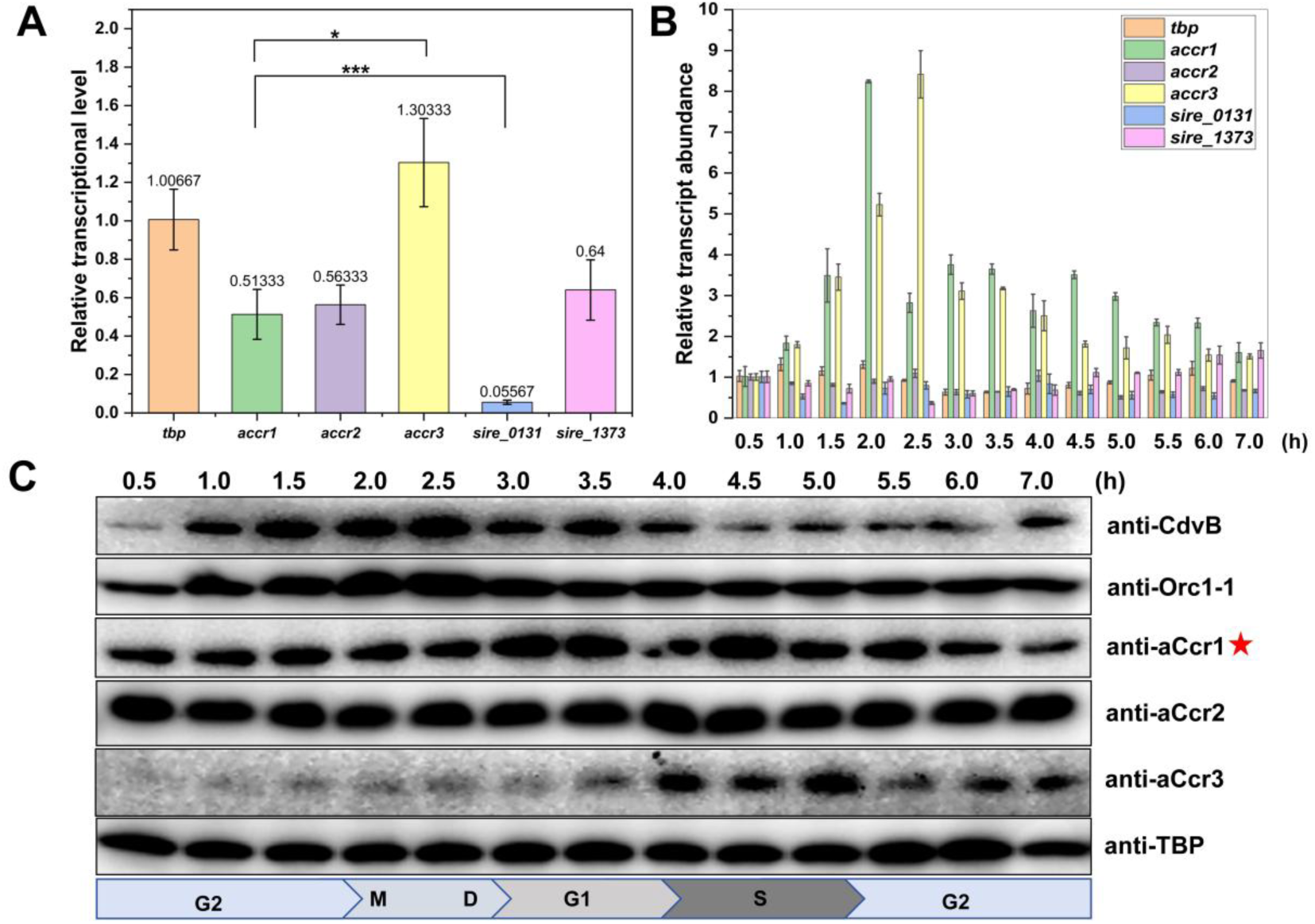
aCcr3 shows cyclic expression at the transcriptional and protein levels, whereas aCcr2 does not. (**A**) RT-qPCR validation of the relative expression of aCcr1 and its homologs in *Saccharolobus islandicus* REY15A, with 16S as an internal reference and *tbp* expression considered to equal 1. Statistical significance was calculated by a One-Sample t-test. *** = p value < 0.0005. ** = p value < 0.005. * = p value < 0.05. (**B**) RT-qPCR validation of the expression of aCcr1 and its homologs during different cell cycle stages, with 16S as an internal reference and *tbp* expression considered to equal 1. (**C**) Western blotting verification of the expression of CdvB, Orc1-1, aCcr1, aCcr2, aCcr3 during different cell cycle phases of the S*accharolobus islandicus* REY15A. TBP was used as a loading control. Notanly, the band of aCcr1 (red star) was not at its predicted position, presumabally due to post-translational modification.

Next, we investigated whether the aCcr1 homologs exhibit cyclic expression. Analysis of the previously sequenced transcriptomes showed that transcription of *accr3* displays a cyclic pattern (Figure S2B) ^24^ and a similar expression pattern was observed in *Sulfolobus acidocaldarius* ^7^. To gain a more refined view on the expression dynamics of RHH domain-containing proteins, the synchronized cultures were sampled every half an hour (instead of 1 hour intervals) and the transcription levels were analyzed using RT-qPCR. Consistent with the transcriptomic data, only *accr1* and *accr3* showed distinct cyclic transcription patterns, whereas the other genes, including *accr2*, were stably expressed (Figure 1B). Notably, the peak transcription of *accr1* occurred approximately 0.5-1 hour earlier than that of *accr3*. Consistently, Western blot analysis showed that aCcr1 protein peaked ∼1 hour earlier than aCcr3 (Figure 1C). Notably, the protein band of aCcr1 migrated at a higher molecular weight than expected, suggesting potential post-translational modifications (PTM). The peak level of aCcr1 expression coincided with cytokinesis, whereas aCcr3 peaked during the G1-S transition (Figure 1C).

### Four of the five aCcr1-like genes are essential for cell viability

In a previous study, we have shown that *accr1* is essential for cell survival ^24^. Of the five *accr1*-like genes, only *sire_0131* could be knocked out (Figure S3A), whereas the genes for aCcr2, aCcr3, and SiRe_1373 all appeared to be essential for cell viability. Notably, *sire_0131* transcripts are present at a very low level (∼5% of the *tbp* expression level) (Figure 1A and S2A), suggesting that SiRe_0131 regulates a single gene or a small number of genes. Furthermore, we observed that there was no significant difference in cell morphology or growth between Δ*sire_0131* and the parental strain E233S (Figure S3B and S3C). These results suggest that SiRe_0131 does not play a measurable role in cell cycle regulation, unlike the other proteins which appear to be critical for the cell survival.

### Overexpression of aCcr2 and aCcr3 resulted in noticeable growth retardation and cell enlargement

To investigate the potential involvement of the essential aCcr1 homologs in cell cycle regulation, we constructed and characterized the overexpression strains of aCcr2 and aCcr3, whereas a strain overexpressing SiRe_1373 could not be obtained (Figure S3D). Overexpression of aCcr2 and aCcr3 led to substantial cell growth retardation, cell enlargement, and an increase in genome copy number, mirroring the phenotypes previously observed during the aCcr1 overexpression (Figure 2A and 2B). Although the overall phenotypes of the aCcr1, aCcr2 and aCcr3 overexpression strains were similar, there were subtle differences between them. Notably, aCcr2 overexpression strains displayed more pronounced growth inhibition compared to aCcr1 overexpression strain, whereas aCcr3 overexpression strain exhibited a moderate level of growth inhibition (Figure 2A). Furthermore, the flow cytometry results showed that 24 hours after arabinose induction, aCcr3-overexpressing cells predominantly contained 2C DNA, whereas the majority of aCcr1- and aCcr2-overexpressing cells were multiploid (Figure 2A).

**Figure 2.**
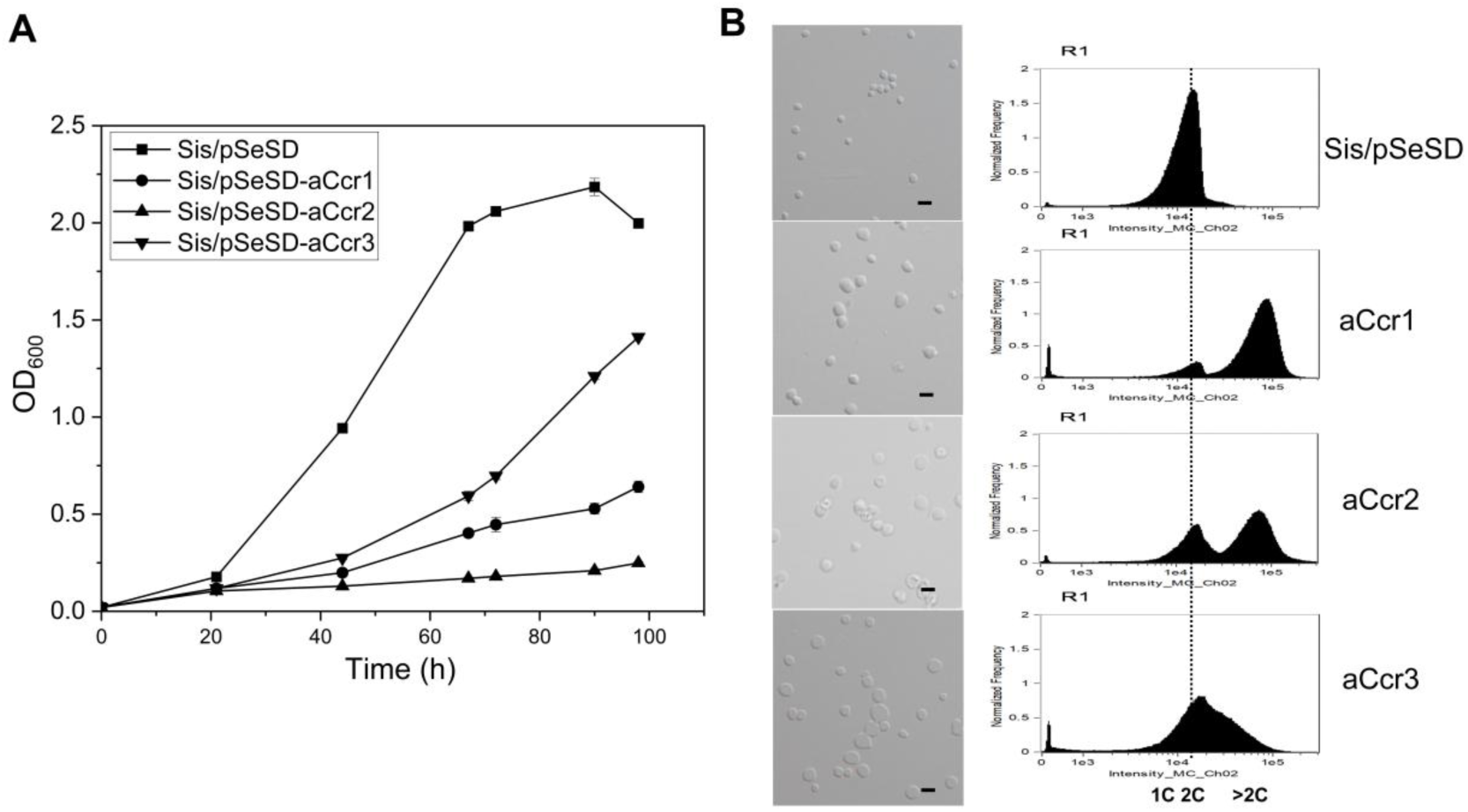
Overexpression of aCcr1, aCcr2 and aCcr3 results in growth retardation and cell enlargement. (**A**) Growth curves of cells over-expressing aCcr1 and its homologs. The cells were inoculated into 30 ml induction medium ATV to a final OD_600_ of 0.03 and the growth was monitored. Error bars represent standard deviations from three independent measurements. Cells harboring the empty plasmid pSeSD were used as a control. (**B**) Bright-field microscopy (DIC) and flow cytometry analyses of cells over-expressing aCcr1 and its homologs. Cells cultured in the induction MTAV medium were sampled at 24h time point for morphological observation and flow cytometry. Scale bars: 2 μm. 1C represents one chromosome, 2C represents two chromosomes, and >2C indicates more than two chromosome copies.

### aCcr1, aCcr2 and aCcr3 have different binding affinities for the *cdvA* promoter

To study the roles of aCcr1 homologs in cell cycle regulation, we analyzed their *in vitro* properties. We successfully expressed and purified all five RHH domain-containing proteins from *E. coli* (Figure S4A and S4B). Analysis of the gel filtration profiles suggested that all proteins form dimers in solution (Figure S4A). Given that the overexpression of aCcr1 homologs (except SiRe_1373) lead to cell enlargement, suggesting an impact on cytokinesis and/or cell cycle, we investigated whether these proteins, similar to aCcr1, exhibit specific binding to the *cdvA* promoter. EMSA experiments with purified proteins showed that aCcr2 and aCcr3 specifically bind to the *cdvA* promoter, whereas SiRe_0131 and SiRe_1373 do not (Figure S4C). These results suggest that *cdvA* is not exclusively regulated by aCcr1, but also by aCcr2 and aCcr3. An overlap in function among aCcr1, aCcr2, and aCcr3 raises a question regarding the essentiality of all three RHH proteins. We hypothesized that the functional coordination of the three aCcr proteins could be achieved through temporal expression and differential binding affinities to the target promoters. Indeed, aCcr1, aCcr2, and aCcr3 displayed differences in their binding affinities to the P*_cdvA_* promoter, with K_D_ for aCcr1, aCcr3, and aCcr2 being 93.49±7.91 nM, 35.95±1.03 nM, and 245.14±7.82 nM, respectively (Figure 3A and 3B). These results indicate that aCcr2 has a much weaker affinity for the *cdvA* promoter compared to aCcr1 and aCcr3.

**Figure 3.**
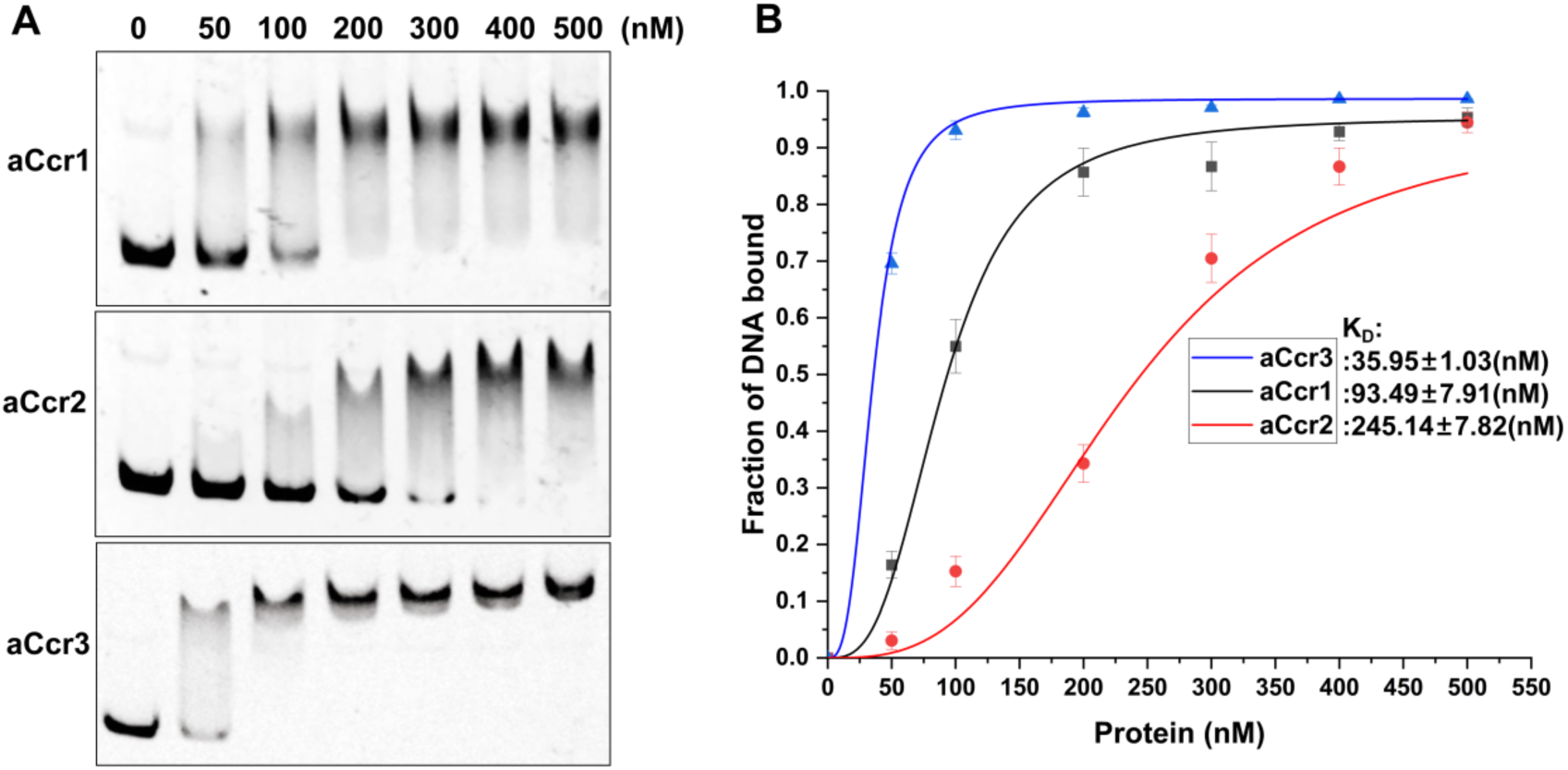
aCcr1, aCcr2 and aCcr3 display differential binding to the *cdvA* promoter. (**A**) EMSA of aCcr1, aCcr2 and aCcr3 binding to the promoter of *cdvA*. The 5’FAM-labelled oligonucleotides were annealed with the respective complementary strands (see Materials and Methods, Supplementary Table S2) and used as the substrate in the EMSA assay. The reaction was performed at 37℃ for 30 min and analysed on a 10% native PAGE. Each reaction contained 2 nM of the FAM-labelled probe and 50, 100, 200, 300, 400 or 500 nM protein. Representative images are shown. (**B**) Quantification of the results in (A). Error bars represent standard deviation from three independent experiments. The data was fitted to logistic curves using Origin 9.

### Many cell cycle-related genes are differentially regulated in the overexpression strains

To identify the target genes regulated by aCcr2 and aCcr3 and to clarify how aCcr1, aCcr2 and aCcr3 coordinate their activities, we conducted transcriptomic analysis of cells overexpressing aCcr2 and aCcr3. The numbers of both upregulated and downregulated (>2-fold change) genes in the aCcr2 and aCcr3 overexpression strains were higher compared to the aCcr1 overexpression strain. Specifically, in the aCcr2 overexpression strain, 437 genes were upregulated and 384 were downregulated, whereas in the aCcr3 overexpression strain, 314 genes were upregulated and 260 were downregulated (Figure 4A and 4B). These numbers are considerably higher than those in the aCcr1 overexpression strain, in which 76 genes were upregulated and 124 genes were downregulated (Supplementary Table S6) ^24^. Notably, 50 and 21 genes were concordantly downregulated and upregulated, respectively, in all three overexpression strains (Figure 4C and 4D). Among the 50 downregulated genes, 10 (20%) were reported to be essential for cell viability (Table 1) ^25^, 24 (48%) exhibited cyclic expression patterns ^24^ and 30 (60%) were in the transcriptionally active chromosome A region (Supplementary Table S3) ^26^, suggesting that aCcr1, aCcr2, and aCcr3 may play a key role in the cell cycle progression by synergistically regulating the transcription of the target genes. Notably, none of the 21 genes that were upregulated were found to be essential (Supplementary Table S4) ^25^. Eight of these twenty-one genes were found to be bound by at least one aCcr1 homolog, 7 genes were expressed cyclically, and 6 were located in the A compartment (Supplementary Table S4). The expression of the non-concordantly regulated genes was specific to aCcr1, aCcr2, or aCcr3. This suggests that, in addition to jointly regulating some of the major cell cycle processes, aCcr proteins also have specialized roles.

**Figure 4.**
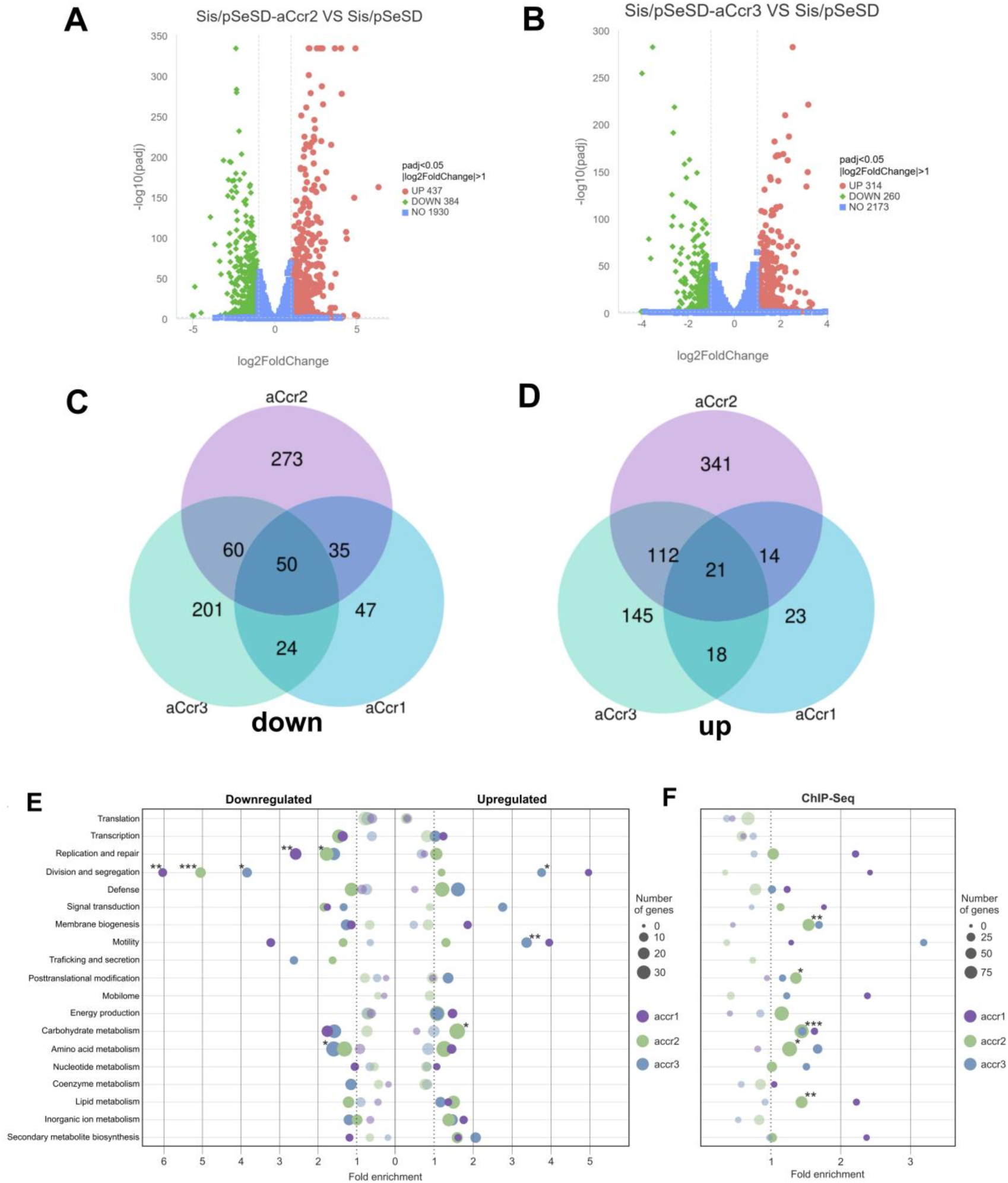
Comparative transcriptomic and functional enrichment analysis of three aCcr proteins. (**A**) and (**B**) Volcano plots of differentially expressed genes in strains Sis/pSeSD-aCcr2 and Sis/pSeSD-aCcr3 compared to the control Sis/pSeSD. X-axis, fold change in gene expression. Y-axis, significance of fold change. Genes exhibiting > 2-fold (i.e. –1 > log2 > +1) significant upregulation and downregulation are highlighted in red and green, respectively, whereas those that showed a <2-fold change in differential gene expression or with no significance are shown in blue. (**C**) and (**D**) Venn diagrams showing the numbers of genes common and differentially downregulated (C) and upregulated by the three aCcr proteins compared with the control. The transcriptomic data for aCcr1 was obtained previously^24^. Functional enrichment analysis of transcriptomics (**E**) and ChIP-seq (**F**) analyses. (E) Dot plot shows the fold enrichment of genes upregulated or downregulated by aCcr1, aCcr2 and aCcr3. Genes were assigned to functional categories provided by the archaeal Clusters of Orthologous Genes (arCOG) ^40^. Genes of unknown function (arCOG categories S and R) were excluded from the analysis. Dot size is proportional to the number of genes up- or down-regulated by each aCcr protein. Statistical significance was calculated using the *enricher* function from the ClusterProfiler package in R ^41^ with Benjamini-Hochberg method correction for multiple comparisons (results can be found in the Supplementary Table 5). (F) Dot plot of the fold enrichment of the aCcr1, aCcr2 and aCcr3 ChIPSeq results. Only peaks found within 1kb of a promoter were considered. Analysis was perfomed as in (E). *** = p value < 0.001; ** = p value < 0.01; * = p value < 0.05.

### Functional enrichment of the differentially regulated genes in the overexpression strains

To analyze which functional categories (as defined in the archaeal clusters of orthologous genes [arCOG]) were statistically significantly overrepresented or underrepresented among the aCcr regulated genes, we performed functional enrichment analysis based on the transcriptomic data (Figure 4E). In the aCcr1 overexpression strain, the significantly downregulated categories included cell division (*cdvA*, *cdvB*, and *cdvC*), chromosome segregation (*segC*), genome replication/repair (the replication-associated genes *orc1-1* and *gins*; the mismatch repair gene *endoMS*), and chromosomal organization (the major chromatin gene *cren7*) (Supplementary Table S5). An even larger fraction of genes implicated in the same functional categories were downregulated in the aCcr2 overexpression strain. Specifically, the downregulated genes in the chromosome segregation category included *segA* and *segB* as well as the gene encoding the chromosome dimer resolution recombinase XerA. The cell division categrory included *cdvA, cdvB1* and *cdvB2*. The aCcr2 overexpression resulted in the downregulation of the same genes in the DNA replication and repair category as in the case of aCcr1, but in addition included genes for reverse gyrase, the replication initiator Whip, DNA polymerase PolB3, and several chromatin-associated proteins (ClsN, aDCP1, SisTrmB1, Sac7d/Sso7d). Concomitantly, aCcr2 overexpression resulted in significant upregulation of the carbohydrate metabolism category, including a total of 35 genes (Supplementary Table S5). The overexpression of aCcr3 led to significant downregulation of functional categories of cell division (*cdvA*, *cdvB* and *cdvC*), chromosome segregation (*SegA*, *SegB,* and *segC* as well as *sire_0136* and *sire_0265* coding for ParA family proteins potentially invovolved in chromosome segregation), and amino acid metabolism (31 genes). The “motility” category was significantly upregulated, with genes for extracellular filaments, including the archaellum locus (*sire_0118*-*sire_0124*). Also upregulated was the putative type IV pilin genes from the bindosome operon (*sire_2517*, *sire_2518*) ^27^, thought to be involved in carbohydrate uptake from the environment ^28^ (Supplementary Table S5). These results suggest that the three aCcr transcription factors repress an overlapping set of key genes involved in cell division, segregation, DNA replication and repair, with additional subsets of genes being aCcr-specific.

### ChIP-seq enrichment of aCcr2 and aCcr3 in exponentially growing cells

To further elucidate the role of aCcr2 and aCcr3 in regulation of specific biological processes *in vivo*, we performed Chip-seq analysis of aCcr2 and aCcr3, and compared the results with those obtained previously for aCcr1 ^24^ (Figure 5A). The analysis revealed the potential aCcr1, aCcr2 and aCcr3 binding sites along the *Sa. islandicus* chromosome, with 610, 1407 and 224 peaks, respectively. For aCcr2, 840 binding sites were enriched within the promoter regions, with 816 within the region between positions -50 and 0 upstream of the start codon, whereas the remaining 567 were within intra- and intergenic regions. For aCcr3, 91 binding sites were identified in the promoter regions, with 89 between -50 and 0, and 133 within intra- or intergenic regions (Supplementary Tables S7 and S8). Motif analysis of the ChIP-seq results for aCcr2 and aCcr3 revealed binding motifs (Figure S5A), similar to that of aCcr1, A(T/G)G(A)TA(G)A(T/G)TACN ^24^. Remarkably, despite the overall similarity in their binding motifs, the binding motifs for aCcr1 and aCcr3 comprise stringent palindromic sequences, whereas that for aCcr2 exhibits greater variability, primarily within the ‘GTA’ sequence. This may explain why aCcr2 can regulate a greater number of genes. Notably, 101 of the ChIP-seq enrichments were shared by all three transcription factors, aCcr1, aCcr2, and aCcr3 (Figure S5B). Notably, the peak enrichment sites were more prevalent in the B compartment (64.3%, 59.2%, and 69.2% for aCcr1, aCcr2, and aCcr3, respectively; Figure S5C). This distribution contrasts with the results from the transcriptomic analysis of cells overexpressing aCcrs, which did not show preferential distribution of the downregulated genes in either A or B compartment, although the upregulated genes were more commonly located in the B compartment (Figure S5D).

**Figure 5.**
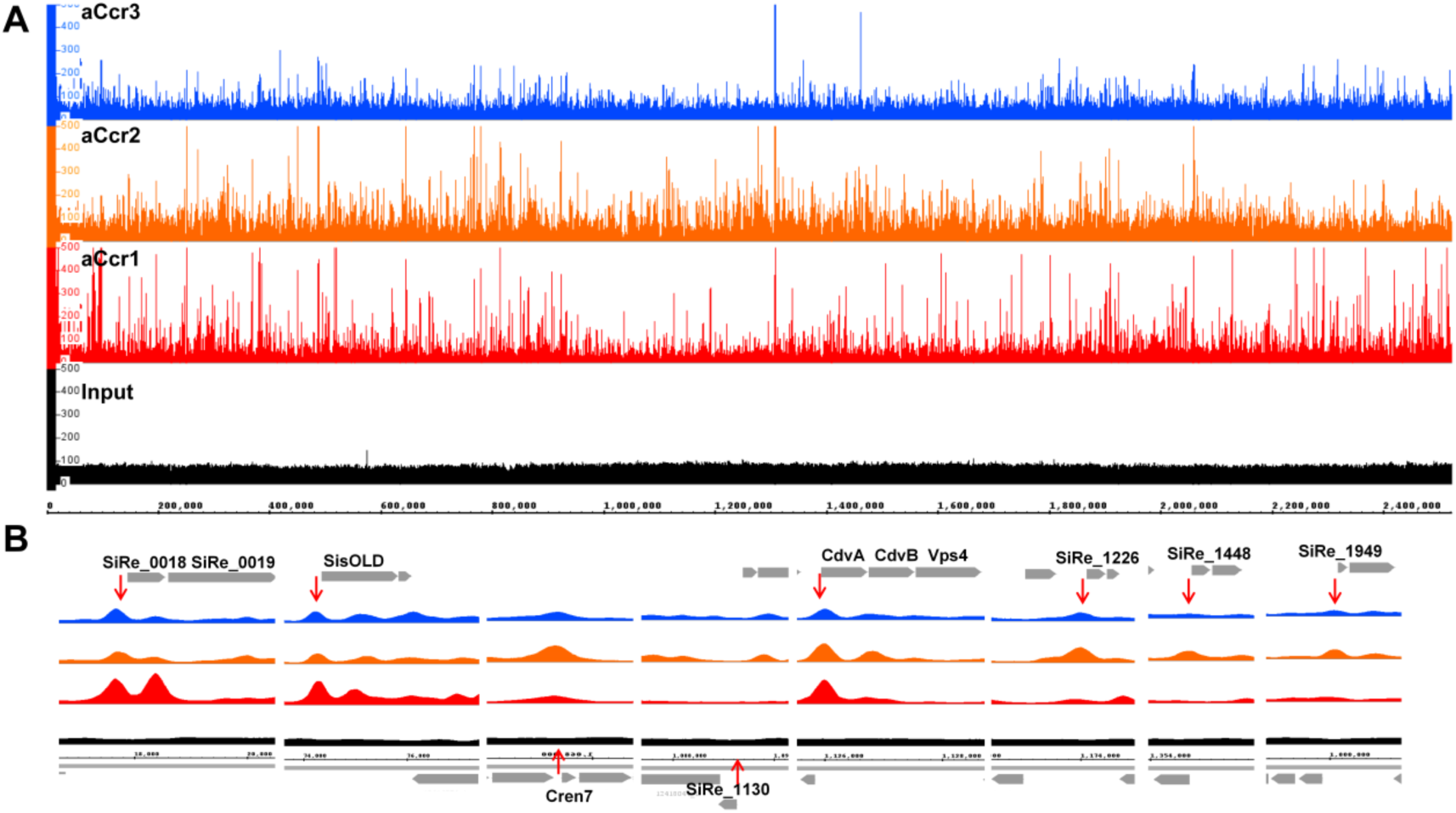
Identification of the aCcr2 and aCcr3 DNA binding sites *in vivo* using Chip-seq. (**A**) Overview of the genomic binding profiles of aCcr1 (red), aCcr2 (orange) and aCcr3 (blue) as monitored by Chip-seq. The sample before immunoprecipitation was used as input. The data on aCcr1 was from our previous report^24^. (**B**) Zoom-in on the promoter regions of ten essential genes co-downregulated by the overexpression of aCcrs.

Functional enrichment analysis of the ChIP-Seq data showed that the aCcr2 binding pattern was enriched in promoters controlling the genes assigned to the categories of membrane biogenesis, posttranslational modification, and carbohydrate, amino acid and lipid metabolisms (Figure 4F). Although these categories were not all significantly enriched in the transcriptomic data (Figure 4E), several genes belonging to these categories were significantly upregulated or downregulated in the aCcr2 overexpression strain (Supplementary Table S6), further highlighting the importance of aCcr2 for a wide range of cellular processes. Overall, both transcriptomic and ChIP-seq analyses suggest that aCcr2 regulates the highest number of genes, possibly due to its constitutive expression throughout the cell cycle and higher tolerance for variations within the promoter motif, whereas aCcr1 and aCcr3 regulate smaller sets of genes that function at specific time points during the cell cycle.

### aCcrs bind to promoters of key cell cycle genes *in vitro*

We have shown that all three aCcr proteins can bind to the promoter of *cdvA* with different affinity (Figure 3). To understand whether other downregulated and ChIP-seq enriched genes can be directly regulated by aCcr proteins, we selected 37 promoters whose genes were the most highly downregulated in the aCcr2-overexpression strain and analyzed whether they can be bound by the three aCcr proteins using the EMSA (Figure S6). The results show that 25 out of 37 (68%) promoter fragments could be bound by at least one of the aCcr proteins. Promoters of 10 genes (*cren7*, *sire_1949*, *segAB*, *xerC*, *orc1-2*, *endoMS*, *ePK1*, *ePK2*, *sire_0018*, and *met*) were bound by all three aCcr protein; promoters of 7 genes (*gins*, *rgy*, *rpo8*, *radA*, *arg*, *cdvB1*, and *sire_1267*) were bound by two aCcr proteins; and those of 8 genes (*whip*, *priL*, *sire_1948*, *sire_1269*, *sire_1448*, *sire_1669*, *sire_*1726 and *sire_1775*) were bound by only one of the aCcrs. The promoters bound by aCcr proteins correspond to the genes that code for proteins involved in many cell cycle processes including chromatin organization (e.g., Cren7, SiRe_1949, SiRe_1948), chromosome segregation and resolution (e. g., SegA, SegB and XerC), DNA replication (e.g., GINS, Whip and Gyrase), DNA repair (e.g., Orc1-2, EndoMS), cell division (CdvA, CdvB1), protein phosphorylation (ePK1 and ePK2), and protein translation (e.g., SiRe_0018 and Met). Only 12 out of the 37 selected promoters (32%) were not bound under our assay conditions by any of the aCcr proteins, suggesting that the expression of these genes is secondarily regulated by the aCcr-controlled gene products. For instance, the chromatin proteins Cren7, SiRe_1949 (Sso7c4 homolog) and SiRe_1948 as well as RHH-domain transcription factor SiRe_0906 are regulated by the aCcr homologs (Supplementary Table S4). Thus, the 12 genes that do not appear to be directly regulated by aCcrs could be controlled through changes in chromatin organization or regulated by transcription factor SiRe_0906. These results provide further evidence that aCcr1, aCcr2, and aCcr3 regulate the archaeal cell cycle progression by transcriptionally controlling the expression of the key cellular functions (see Supplementary Notes for detailed description of the aCcr-affected processes).

## Discussion

Recently, it has been shown that expression of many of the *Sa. islandicus* genes is coordinated with different phases of the cell cycle, with the overall logic being similar to that in certain eukaryotes, such as *Saccharomyces cerevisiae* ^21^. However, the regulation of the cell cycle progression remained unclear. Here, we have shown that aCcr1, aCcr2 and aCcr3 regulate the expression of an overlapping array of genes involved in many key cellur processes (Figure 4E) and propose that the cell cycle regulatory network of *Sa. islandicus* is centered around the aCcr proteins (Figure 6). aCcr1 and aCcr3 act as repressors during specific time points, namely, the M/G1 and G1/S transitions, respectively, which ultimately determine the progression of the cell cycle. By contrast, aCcr2 functions as a more general repressor, acting throughout the cycle. Specifically, aCcr1 represses the expression of genes involved in chromosome segregation, cell division, and phosphorylation during a late stage of cytokinesis. aCcr3, which is expressed during the G1-S phase, represses the expression of genes involved in DNA replication, DNA repair, RNA turnover, and amino acid metabolism. By contrast, continuous expression of aCcr2 throughout the cycle controls a wider range of genes, including repression of chromatin genes and activation of genes related to extracellular appendages and carbohydrate metabolism.

**Figure 6.**
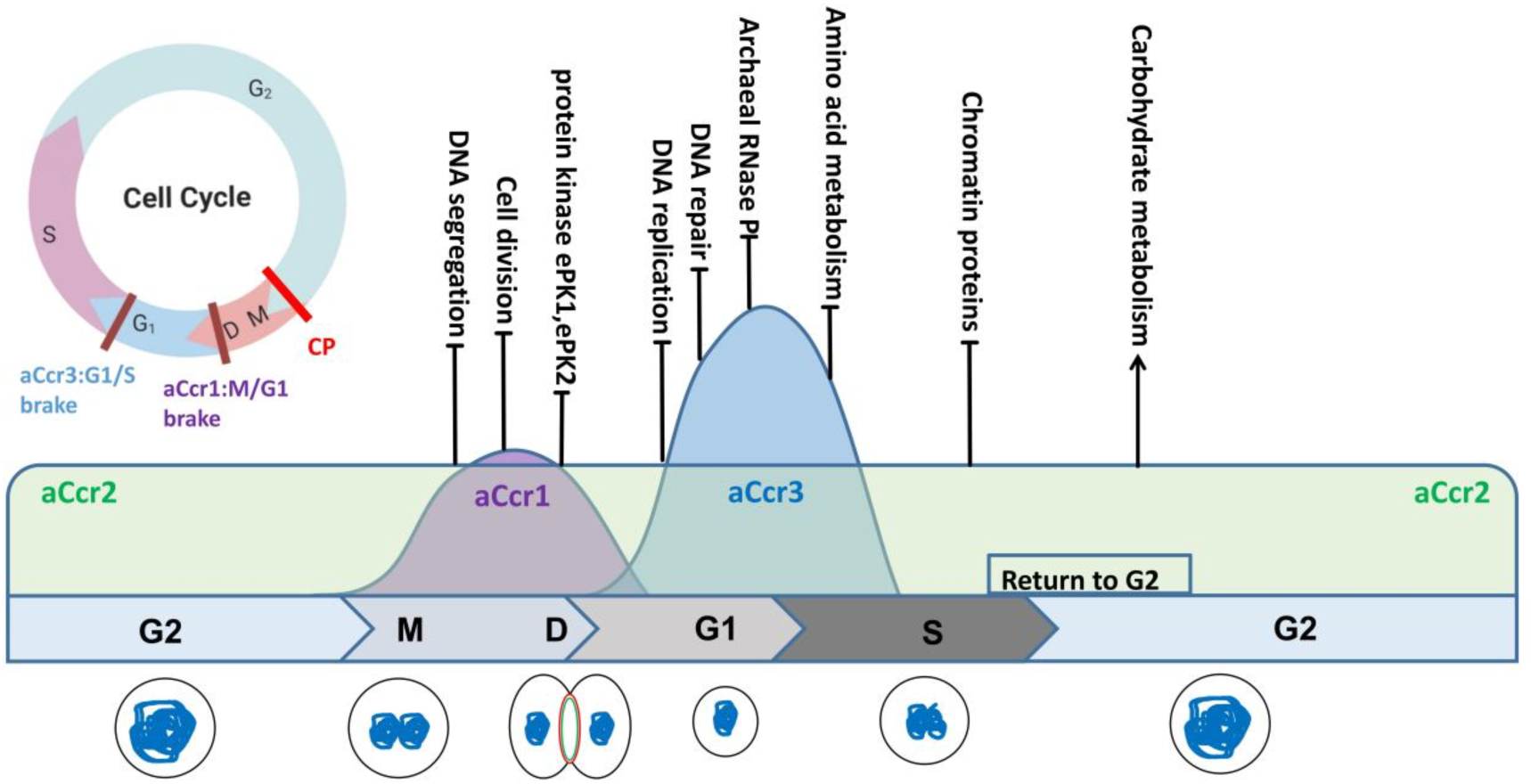
Model of cell cycle regulation by aCcrs in Sulfolobales. Cell cycle progression is controlled by aCcrs at different stages. Specifically, aCcr1 regulates the expression of genes involved in chromosome segregation, cell division, and phosphorylation during late stage of cell division. aCcr3, which is highly expressed during G1-S phase, represses the expression of genes involved in DNA replication, DNA repair, RNA turnover, and amino acid metabolism. aCcr2 is continuously expressed throughout the cycle, controlling the expression of a wider range of genes, including chromatin and carbohydrate metabolism. Specific biological processes suppressed or activated (red arrow) by aCcrs during specific stages of *Saccharolobus islandicus* cell cycle are shown. CP: G2 checkpoint, which was potulated in a recent report^29^. We define the function of aCcr1 and aCcr3 in the regulation network as M/G1 and G1/S braking points, respectively.

Eukaryotic cells have evolved three main checkpoints that ensure normal progression of the cell cycle. These include the G1 checkpoint, G2/M checkpoint and the spindle checkpoint ^10,11^. Whether checkpoint-like control of the cell cycle progression also operates in archaea, such as Sulfolobales, remains poorly understood, but recent evidence suggests in favour of an analogous mechanism. The fact that Sulfolobales cells can be synchronized to the late G2 phase using acetic acid suggests the existence of a cell division entry commitment point, a putative G2/M checkpoint. Notably, similar to certain eukaryotic viruses, several archaeal viruses arrest cell division, further highlighting the G2/M transition as a critical point of the cell cycle progression ^13^. A recent study has shown that the correct assembly of the cell division ring is prerequisite for the initiation of genome condensation into two spatially separated nucleoids, a regulatory mechanism resembling the eukaryotic spindle assembly checkpoint^29^. The cytokinesis itself is controlled by the proteasome which degrades the major divisome component CdvB ^14,16^, with the cyclically-expressed ArsR-type repressor, CCTF1, controlling the timely expression of the proteasome-activating nucleotidase (PAN) ^30^. Furthermore, we have recently shown that phosphorylation of the α subunit of the proteasome by a cyclically expressed kinase (ePK2) inhibits the proteasome assembly and regulates the degradation of the cell division proteins CdvB and CdvB1 in *Sa. islandicus* ^31^. Collectively, these results highlight the multi-layered control of cell cycle in Sulfolobales immediately prior and during the cytokinesis, operating at the transcriptional and post-translational levels. By contrast, aCcr1 and aCcr3 in *Sa. islandicus* appear to control the exit from cytokinesis and transition between the G1 and S phases, respectively.

Strict cell cycle regulation allows only one cell division and one genome duplication to occur during a single cell cycle. Based on the expression timing, we propose that following a successful round of chromosome segregation and initiation of cytokinesis, the major role of aCcr1 is to repress the genes implicated in chromosome segregation and cell division, safeguarding the progression into G1. By contrast, aCcr3 is expressed during the transition between the G1 and S phases. Notably, the expression of most genome replication proteins takes place during the G1 phase ^21^. Thus, the primary role of aCcr3 could be repression of the replisome genes (see Supplementary notes) to prevent the reinitiation of genome replication. Indeed, the profile of DNA content analyzed by flow cytometry in the aCcr3 overexpression strain was radically different from those of aCcr1 and aCcr2 overexpression strains. In particular, the majority of the aCcr3-overexpressing cells contained two chromosome copies, whereas overexpression of either aCcr1 or aCcr2 resulted in accumulation of polyploid cells (Fig. 2B), suggesting that cell division but not genome replication was blocked in the latter strains. aCcr3 has the highest affinity for the shared promoters among the three aCcr transcription factors, suggesting that it can displace its homologs when expressed. The affinity of aCcr2 for binding to the promoters of the shared genes, such as *cdvA*, is the weakest, suggesting that it can be displaced in a temporal fashion by transcription activators (prior cytokinesis) and aCcr1 or aCcr3 (following cytokinesis). Alternatively or additionally, PTMs may play a role in the promoter ‘handover’ between aCcr1, aCcr2, and aCcr3. This possibility is supported by our observation that aCcr1 appeared to be posttranslationally modifed (Fig. 1C). Notably, aCcr1 homologs contain several conserved serine and threonine residues (Fig. S1A) and phosphorylation at the sixth threonine of aCcr2 has been reported in *Sulfolobus acidocaldarious^32^*. This threonine is proximimal to the conserved DNA binding lysine residue; thus, Thr6 phosphorylation by a cyclically expressed kinase, such as ePK2, could lower the promoter biding affinity during strategic cell cycle time points. Thus, the orderly progression of the Sulfolobales cell cycle appears to be controlled by a set of repressors which inhibit the expression of key cellular processes during strategic time points, which we refer to as the “braking points”.

How gene expression is activated, especially in the case of aCcrs themselves, remains unclear. It has been suggested that TFB2 (SiRe_1144), which has the cyclin fold and is cyclically expressed in *Sa. islandicus*, serves as a potential transcriptional activator ^22^, but more experimental evidence is needed to confirm this hypothesis. Furthermore, analysis of ATAC-seq data from *Sa. islandicus* REY15A has revealed chromatin-free regions (CFRs) around the promoters that encompass all core promoter elements, such as BRE, TATA, and regions upstream of the transcription start site ^33,34^. Interestingly, ATAC-seq data from the halophilic euryarchaeon *Haloferax* has shown that there is no apparent correlation between promoter accessibility and gene expression, suggesting that its promoters may be in a constitutively accessible state ^34^. It is thus possible that expression of the key cellular processes in Sulfolobales is primarily regulated by repressors, rather than specific activators, and aCcr homologs could be some of the key players for the coordination of the cell cycle. We anticipate that further research on Sulfolobles and other archaea will provide additional insights into the evolution of the eukaryotic cell cycle regulation systems.

## Materials and Methods

### Strains, growth conditions and transformation of the strains

*Saccharolobus islandicus* REY15A was aerobically grown with shaking at 150 rpm at 75 ℃ in MSTV medium, which contained mineral salts (M), 0.2% (wt/vol) sucrose (S), 0.2% (wt/vol) tryptone (T), and a mixed vitamin solution (200×V). The mutant strain E233S (REY15AΔ*pyrEF*Δ*lacS*) was cultivated in STVU medium with an additional 0.01% (wt/vol) uracil (U). The pH of the culture was adjusted to 3.3 using sulfuric acid, following previously published methods ^13,24^. MSTV medium was utilized for selecting uracil prototrophic transformants. Culture plates were prepared using gelrite (0.8% [w/v]) by combining 1 × MSTV with an equal volume of 1.6% gelrite. MATV medium containing 0.2% (wt/vol) arabinose (A) was employed for inducing protein overexpression. The plasmids and strains employed in this study are listed in Tables S1 and S2, respectively.

### Bright-field microscopy

For bright-field microscopy analysis, 5 μl of cell suspension at specified time points were observed using a NIKON TI-E inverted fluorescence microscope (Nikon, Japan) in differential interference contrast (DIC) mode.

### Cell cycle synchronization

The *Sa. islandicus* E233S strains were synchronized following established protocols ^35^. Initially, the cells were grown aerobically at 75℃ with shaking at 145 rpm in 30 ml of STVU medium. Once the OD_600_ reached 0.6-0.8, the cells were transferred to 100 ml STVU medium with an initial estimated OD_600_ of 0.05 and cultured under the same conditions. Upon reaching an OD_600_ of 0.15-0.2, acetic acid was introduced at a final concentration of 6 mM to arrest the cells at the G2 phase of the cell cycle after 6 hours of treatment. Subsequently, the cells were harvested by centrifugation at 3,000 g for 10 minutes at room temperature to eliminate the acetic acid, followed by two washes with 0.7% (w/v) sucrose. The cells were then resuspended in 100 ml of pre-warmed STVU medium and cultured as previously described for further analysis. Flow cytometry was used to analyze the cell cycle, with samples collected at specific time points for subsequent Western blotting or RT-qPCR analysis.

### Flow cytometry analysis

The cell cycle of synchronized E233S cells and strains overexpressing aCcr1 homologs was analyzed using flow cytometry with an ImageStreamX MarkII quantitative imaging analysis flow cytometer from Merk Millipore, Germany. Cell samples were fixed with 70% ethanol for at least 12 hours at each time point and stained with propidium iodide (PI) at a concentration of 50 mg/ml. Data was collected for a minimum of 500,000 cells per sample and analyzed using IDEAS data analysis software.

### Protein expression and purification

Recombinant aCcr1 and its homologs with a C-terminal His-tag were expressed in *Escherichia coli* BL21(DE3)-RIL cells. The protein expression was induced during the logarithmic growth phase (OD_600_=0.4∼0.8) by adding 0.5 mM isopropyl-thiogalactopyranoside (IPTG), followed by cultivation at 37 ℃ for 4 hours. The cells were then collected by centrifugation, and the cell pellet was resuspended in buffer A (50 mM Tris-HCl pH 8.0, 300 mM NaCl, 5% Glycerol). After lysing the cells by sonication, the cell extract was clarified by centrifugation at 13,000 × *g* for 20 minutes at 4℃. The supernatant was incubated at 70℃ for 20 minutes, centrifuged again at 12,000 *g* for 20 minutes, and filtered through a 0.45 μm membrane filter. The samples were then loaded onto a Ni-NTA agarose column (Invitrogen) pre-equilibrated with buffer A and eluted with a linear imidazole gradient (40-300 mM imidazole) in buffer A. The fractions were pooled, concentrated to 1 ml, and further purified by size exclusion chromatography (SEC) using a Superdex 200 column with buffer A. Finally, the samples were dialyzed in a storage buffer containing 50 mM Tris-HCl pH 7.4, 100 mM NaCl, 1mM DTT, 0.1 mM EDTA, and 50% glycerol. The protein concentrations were determined using the Bradford protein assay kit (Beyotime), and the purity was analyzed by 15% SDS-PAGE stained with Coomassie blue.

### Western blotting

The expression levels of aCcr1 and homologous proteins in synchronized cells were analyzed using Western blotting. Approximately 2×10^8^ cells were collected at specific time points for each sample and subjected to SDS-PAGE analysis on a 15% gel. The separated proteins were then transferred onto a PVDF membrane. Specific bands were detected using chemiluminescence with Kermey ECL Super Western Blotting Detection Reagents (KermeyBiotech, Zhengzhou, China) as per the manufacturer’s instructions. Primary antibodies against CdvB, aCcr1, and TBP were generated in rabbits by HuaAn Biotechnology Co., Ltd. (Hangzhou, Zhejiang, China)^35,36^, whereas the Orc1-1 antibody was from the laboratory stock of Prof. Qunxin She^37^. Antibodies against aCcr2 and aCcr3 were produced by AtaGenix Laboratories Co., Ltd. (Wuhan), using purified recombinant proteins from E. coli in rabbits. The primary antibody against the His-tag was purchased from TransGen Biotech Company (Beijing, China). Goat anti-rabbit antibodies (KermeyBiotech, Zhengzhou, China) conjugated with peroxidase were utilized as secondary antibodies.

### Electrophoretic mobility shift assay (EMSA)

The binding capacity of aCcr1 and its homologs proteins was assessed using a DNA substrate consisting of a 100 nucleotide (nt) strand labeled with 5’-FAM at the *cdvA* gene and other gene promoters (Supplementary Table S2). The DNA substrates were amplified using specific primers and cloned into pUC19 plasmids at EcoRI and HindIII sites, resulting in a series of pUC19 plasmids with different promoters. FAM-labeled substrates were generated by PCR using FAM-pUC19-EcoRI-F/Specific gene promoter-R primers and various pUC19 plasmids as templates ^38^. For the binding assay, a 20 μl reaction mixture containing 10 nM dsDNA, 50 mM Tris-HCl (pH 7.4), 5 mM MgCl_2_, 20 mM NaCl, 50 μg/mL BSA, 1 mM DTT, 5% glycerol, and varying concentrations of purified proteins was prepared. The reaction mixture was incubated at 37 ℃ for 30 minutes before being loaded onto a 12% native polyacrylamide gel. Following electrophoresis in 0.5×TBE buffer, the gel was visualized using an Amersham ImageQuant 800 biomolecular imager (Cytiva).

### Transcriptome analysis

Strains of Sis/pSeSD and Sis/pSeSD-aCcr2, Sis/pSeSD-aCcr3 were cultured in ATV medium following the described conditions. For transcriptomic analysis, the culture was inoculated with an initial OD_600_ of 0.1. After 12 hours of cultivation, the cells were pelleted at 6,000 × *g* for 10 minutes. The pellet was then resuspended in 1 ml of PBS buffer, pelleted again, and stored at -80℃. Total RNA was extracted using the Trizol reagent (Ambion, Austin, TX, USA) and the RNA Nano 6000 Assay Kit of the Bioanalyzer 2100 system (Agilent Technologies, CA, USA) was used to assess the total amounts and integrity of the RNA. Transcriptomic analysis was carried out by Novogene (Beijing, China), with approximately 3 μg of high-quality RNA per sample used for the construction of RNA-Seq libraries. To begin, mRNA was purified from total RNA using probes to remove rRNA. First strand cDNA was then synthesized using a random hexamer primer and M-MuLV Reverse Transcriptase, followed by RNaseH treatment to degrade the RNA. In the DNA polymerase I system, dUTP was utilized to replace dTTP in dNTP to synthesize the second strand of cDNA. Exonuclease/polymerase activities were used to convert remaining overhangs into blunt ends, followed by adenylation of the 3’ ends of DNA fragments. Adaptors with hairpin loop structures were ligated to prepare for hybridization. Finally, the USER Enzyme was employed to degrade the second strand of cDNA containing U. To select cDNA fragments of a desired length range of 370∼420 bp, the library fragments were purified using the AMPure XP system from Beckman Coulter in Beverly, USA. Following PCR amplification, the product was further purified with AMPure XP beads to obtain the final library. Subsequently, the libraries were sequenced utilizing the Illumina NovaSeq 6000 platform. The clean reads obtained were then aligned to the reference genome sequence of *Sa. islandicus* REY15A^39^. The resulting data was subjected to Fragments Per Kilobase of transcript sequence per Million base pairs sequenced (FPKM) analysis to ascertain the expression levels of all genes within the *Sa. islandicus* genome. Differential expression analysis of the genome (comparing over-expression of aCcr2 or aCcr3 versus an empty vector) was carried out using the DEGSeq R package. To ensure statistical reliability, the resulting P-values were adjusted using the Benjamini and Hochberg’s method for controlling the false discovery rate. A threshold of padj<0.05 and |log2(foldchange)| > 1 was set to determine significantly differential expression.

### Functional enrichment analysis

Genes were assigned to functional categories provided by the archaeal Clusters of Orthologous Genes (arCOG)^40^. Genes of unknown function (arCOG categories S and R) were excluded from the analysis. Statistical significance was calculated using the enricher function from the ClusterProfiler package in R^41^ with Benjamini-Hochberg method correction for multiple comparisons.

### Quantitative reverse transcription PCR (RT-qPCR)

Quantitative reverse transcription PCR (RT-qPCR) was utilized to analyze changes in the expression levels of aCcr1 and its homologous genes. Samples were collected from E233S strains during the logarithmic growth phase and after synchronization at the corresponding time points. Total RNA extraction was carried out using SparkZol (SparkJade Co., Shandong, China). First-strand cDNAs were synthesized from the total RNA following the protocol of the First Strand cDNA Synthesis Kit (Accurate Biotechnology Co., Hunan, China) for RT-qPCR analysis. The resulting cDNA samples were used to measure the mRNA levels of the target genes through qPCR using the SYBR Green Premix Pro Taq HS qPCR Kit (Accurate Biotechnology Co., Hunan, China) and gene-specific primers (Supplementary Table S2). PCR was conducted in a CFX96TM (Bio-Rad) instrument with the following steps: denaturation at 95℃ for 30 seconds, followed by 40 cycles of 95℃ for 5 seconds and 60℃ for 30 seconds. The relative quantities of mRNAs were determined using the comparative Ct method with the 16S rRNA gene as the internal reference. All qPCR primers were verified to have an amplification efficiency within the range of 90% to 110% (Figure S1C).

### Chromatin immunoprecipitation (ChIP-seq)

Chromatin immunoprecipitation (ChIP-seq) was conducted following the protocol outlined by Blombach. et al^42^. with slight adjustments. In brief, 500 ml of E233S strains were harvested during the exponential growth phase and cross-linked with 0.4% formaldehyde solution. The cross-linking process took place for 5 minutes and was halted by the addition of Tris/HCl pH 8.0 to a final concentration of 100 mM. Subsequently, the samples were cooled on ice for 5 minutes and then centrifuged (6,000 × *g*, 20 minutes, 4 ℃). The cells were resuspended in 10 mL of PBS, pelleted using an Eppendorf Centrifuge 5410 R (6,000 *g*, 20 minutes, 4 ℃), and washed with PBS. The pelleted cells were cryopreserved in liquid nitrogen and stored at -80 ℃. Next, the cells were resuspended in 3ml of Lysis buffer (50 mM HEPES-NaOH pH 7.5, 140 mM NaCl, 1mM Na_2_EDTA, 0.1% sodium deoxycholate, 1% TritonX-100) and sonicated to fragment the DNA to a size range of 200-500 bp. After centrifugation (10,000 × g for 15 minutes), a 50 µl aliquot of the DNA-containing supernatant was preserved as an input control, while the remaining sample was divided into 200 µl aliquots. Each 200 µl aliquot was incubated with 3mg of anti-aCcr2/aCcr3 antibody at 4 ℃ overnight. The following day, 50 μL of resuspended protein A beads (Cytiva) were added to each ChIP assay, and the tubes were rotated at 4℃ for 2 hours. After harvesting the beads, the supernatant was decanted. The immune complexes were isolated through centrifugation and subjected to three consecutive 15 minutes incubations with 1 ml of Lysis buffer, each with vigorous shaking at 4°C. The beads were then washed once with Lysis buffer containing 500 mM NaCl, followed by another wash with LiCl wash buffer consisting of 10 mM Tris-HCl pH 8.0, 100 mM LiCl, 1 mM Na_2_ EDTA, 0.5% sodium deoxycholate, and 0.5% Nonidet P-40. The beads were resuspended in 1 mL of TE buffer (10 mM Tris-HCl pH 8.0, 1 mM Na_2_ EDTA) and incubated for 15 minutes at 4℃. Subsequently, the immune complexes were disrupted by resuspending the beads in Chip-Elute buffer containing 50 mM Tris (pH 8.0), 10 mM EDTA, and 1% SDS, and heating the sample at 65℃ for 30 minutes. The beads were then removed, and the DNA was extracted by treating the samples with 2 μL of 10 mg/mL RNase A and 5 μL of 20 mg/mL proteinase K overnight at 65℃. Following incubation, the samples were collected and the captured DNA was purified using the DNA Cycle-Pure Kit (Omega) according to the manufacturer’s instructions. The input samples underwent the same treatment procedure as described above, excluding the addition of antiserum and beads. Two biological replicates were carried out for aCcr2 and aCcr3, respectively. The purified DNA was utilized for ChIP-seq library preparation. The library was constructed by Novogene Corporation (Beijing, China), followed by pair-end sequencing of the sample on an Illumina platform (Illumina, CA, USA). Library quality was evaluated using the Agilent Bioanalyzer 2100 system, with clean data obtained by removing low-quality reads, as well as reads containing adapters and poly-N from the raw dataset. All subsequent analyses were conducted using the clean, high-quality reads. Reference genome and gene model annotation files were sourced from GenBank, and the reads were aligned to the REY15A genome ^39^. The entire genome was scanned with a specific window size using MACS2 (version 2.1.0)^43^ to calculate the read enrichment level and identify IP enrichment regions for peak calling. A q-value threshold of 0.05 was applied to all datasets. Following peak calling, visualizations were created to display the distribution along the chromosome, peak width, fold enrichment, significance level, and peak summit number per peak. Peaks located at -100 to 0 of the genes were extracted and analyzed. Lastly, the MEME-ChIP tool^44^ was utilized to discover motifs within the peak regions.

## DATA AVAILABILITY

All data supporting the findings of this study are available within the article and its Supplementary Information, or from the corresponding author upon reasonable request.

## ACKNOWLEDGEMENTS

We would like to thank members of the CRISPR and Archaea Biology Research Center for helpful discussions and technicians from the Core Facilities for Life and Environmental Sciences, State Key Laboratory of Microbial Technology of Shandong University for assistance. This work was supported by National Natural Science Foundation of China [32393973 and 32370033] to Y.S. and Postdoctoral fellowship Program of CPSF (GZC20231471) and Postdoc Innovation Project of Shandong Province (SDCX-ZG-202400122) to Y.Y.

## Author contributions

**Yunfeng Yang**: Conceptualization; Data curation; Formal analysis; Investigation; Visualization; Methodology; Writing-original draft and editing. **Shikuan Liang**: Investigation; Methodology. **Zixin Geng**: Investigation; Methodology. **Miguel V. Gomez-Raya-Vilanova**: Functional enrichment analysis; Investigation; Writing-review and editing. **Wenying Xia**: Investigation; Methodology. **Junfeng Liu**: Methodology. **Qihong Huang**: Methodology. **Jinfeng Ni**: Writing-review and editing. **Qunxin She**: Conceptualization; Writing-review and editing. **Mart Krupovic**: Conceptualization; Formal analysis; Writing-review and editing. **Yulong Shen**: Conceptualization; Supervision; Funding acquisition; Project administration; Writing-review and editing.

