## Supplementary figures and tables for "Three transcription factors coordinate the archaeal cell cycle progression through a regulatory braking point mechanism"

### SUPPLEMENTARY NOTES

#### Major cell cycle processes are regulated directly or indirectly by aCcr proteins

*Chromosome dimer resolution, segregation, and cell division (G2-M/G1)*

Following genome replication, chromosomal dimers can be formed and have to be resolved by the XerA recombinase during the G2 phase, prior to segregation during the M phase [1,45]. Our transcriptomics analyses revealed that *xerA* was down-regulated in the aCcr2-overexpressing cells. Consistently, the EMSA analysis showed that all three aCcr proteins can directly bind to the $P_{xerA}$ (*sire_1625*; Figure S6). The chromosome segregation-associated genes *segAB* were significantly down-regulated in the aCcr2 overexpressing strain, and their promoter region contains an aCcr-box motif (AGTATTAT), indicating that *segAB* transcription is also under the control of aCcrs (Figure S6). Additionally, genes encoding a ParA-family ATPase (SiRe_0265) and the chromosome segregation-associated protein SegC[46] (SiRe_1963) were both downregulated in the aCcr overexpression strains (Table S3), implying that the process of chromosome segregation in *Sa. islandicus* is under strict regulation and that SiRe_0265 may be an additional component of the chromosome segregation of *Sa. islandicus*. Consistent with the *in vitro* binding experiments (Figure S4C), *cdvA* was down-regulated in aCcr1, aCcr2, and aCcr3 overexpression strains (Table 1). Additionally, another cytokinetic gene, *vps4*, was also down-regulated in all three overexpression strains (Table 1). Given that *vps4* (*cdvC*) and *cdvB* form a bicistronic operon [47], expression of both genes is likely affected simultaneously. Notably, EMSA experiments showed that the *cdvB1* promoter could be apparently be bound more tightly by aCcr3 than aCcr1 (Figure S6). In summary, we demonstrated that aCcrs regulates chromosome dimer resolution, segregation, and cell division from promoter sequence analysis and EMSA experiments.

*DNA replication, DNA damage response, and repair (M/G1-S)*

The replication initiation protein genes *orc1-1* (Figure 1C), *orc1-3*, and *whip* exhibit high expression levels during the M/G1 phase [21], suggesting that replication initiation proteins are prepared prior to the replicative S phase. Despite the repression of *orc1-1* in the three overexpression strains, aCcrs did not bind to the corresponding promoters (Fig. S6). We hypothesized that Orc1-1 expression might be indirectly regulated by aCcrs. By contrast, the promoter of *whip*, $P_{whip}$, was bound by aCcr1 (Fig. S6), consistent with the presence of the aCcr-box-like palindromic sequence (AGTAGTAC) in the corresponding promoter. Coordinated expression of the replication initiators orchestrates genome replication, preventing multiple rounds of replication prior to cell division. Additionally, the *gins* gene coding for a subunit of the replication initiation complex CMG (Cdc45-MCM-Gins) is also repressed by aCcr1 and aCcr3, further ensuring accurate coordination of genome replication. *Gins* (*sso0772/sire_1229*) and *mcm* (*sso0774/sire_1228*) form an operon [47], suggesting that aCcr1 and aCcr3 regulate the expression of both genes. The primase complex PriSLX is responsible for primer synthesis during chromosome replication in *Sa. islandicus*. The transcription of the regulatory PriX and the large PriL subunits is regulated by aCcr3 (Supplementary Table S5). Additionally, the gene for the PolB1-binding proteins PBP1 (SiRe_1861) was down-regulated by the expression of aCcrs (Supplementary Table S3), although its cyclic expression is not clearly defined. *sire_1949* encodes a homolog of Sso7c4, a predicted chromatin protein, and is another essential genes directly regulated by aCcrs (Table 1 and Figure S6). The protein is highly expressed during the S and G2 phases, suggesting a role in genome replication or maintenance. Reverse gyrase, a type I topoisomerase, which uses ATP hydrolysis to introduce positive supercoiling into DNA, plays a crucial role in replication, recombination, and other genetic processes [48,49]. Transcriptomic data and EMSA experiments have revealed that reverse gyrase gene (*sire_1124*) is directly regulated by aCcrs (Supplementary Table S6 and Figure S6).

DNA damages may occur during the cell cycle and are repaired immediately or presumably at the the G2 phase. Orc1-2 plays a central role in the DNA damage response in *Sa. islandicus* [15], but its activation mechanism remained unclear. Here we show that the *orc1-2* promoter, which contains the aCcrs-box motif GGTAATAC, is bound by aCcrs (Figure S6), potentially explaining the low expression of *orc1-2* in the absence of DNA damage. Upon exposure to damaging agents, *orc1-2* expression increases, leading to activation of the downstream genes. Transcriptomic analysis of cells treated with the DNA damaging agent NQO revealed downregulation of both aCcr1 and aCcr3 [15]. We hypothesize that the observed decrease in the aCcr1 and aCcr3 levels lifts the *orc1-2* transcriptional repression, thereby activating the DNA damage response. It was proposed that EndoMS, the archaea-specific DNA mismath repair nucelase (SiRe_0025), is involed in mismatch repair at the replication fork through homologous recombination, in which helicase (SiRe_0017) and RecA/RadA recombinases (SiRe_1224 and SiRe_1747) are also involved [50]. The promoter for EndoMS was bound by aCcrs (Figure S6). And genes related to repair, such as those coding for the HerA helicase (SiRe_0017), RecA/RadA recombinases (SiRe_1224 and SiRe_1747), and Lhr-like helicase (SiRe_1605), are also downregulated in strains overexpressing aCcrs (Table S3). Taken together, our data strongly suggest that aCcrs regulate genome replication, repair, and DNA damage response.

*Amino acid and tRNA metabolisms, and protein synthesis (M/G1-S)*

Among the essential genes directly regulated by aCcrs, we discovered two genes located in the same operon, *sire_0018* and *sire_0019* (Figure 5B and Table 1). Queuine tRNA-ribosyltransferase, encoded by *sire_0018* and *sire_0019*, plays a crucial role in tRNA synthesis by transferring Queuine, a unique nucleoside, to specific tRNA molecules[51]. This process is vital for optimizing translation rates and maintaining proteome stability. Additionally, aCcrs regulate a number of other genes involved in translation. These include genes encoding several tRNA-related enzymes (SiRe_1238, SiRe_1973, SiRe_0420, and SiRe_2216), an amino acid transporter (SiRe_0688), subunits of RNase P (subunit p30, SiRe_1269; subunit p14, SiRe_1270), a ribonucleoprotein enzyme that cleaves precursor sequences from the 5' ends of pre-tRNAs [52], certain tRNAs (tRNA-Met and tRNA-Arg; Supplementary Table S6 and Figure S6), exosome subunit (SiRe_1268), and ribosomal protein L15E (SiRe_1267). Consistently, an aCcr-binding motif of GTAATAC was found in the promoter region of these genes. Notably, the initiation codon Met plays a crucial role in protein synthesis, impacting the entire translation process. Thus, transcriptional control of the tRNA-Met promoter by aCcrs suggests a prominent control over the translation process.

*Eukaryotic-like kinases (M/G1-S)*

Although Sulfolobales archaea lack homologs of eukaryotic cyclins, they encode analogs of eukaryotic protein kinases (ePKs). SiRe_2056 (ePK1) has been shown to primarily perform protein phosphorylation in response to various cellular stresses in *Sa. islandicus* REY15A [53]. The function of another eukaryotic analogous kinase, SiRe_2030 (ePK2), is less well understood in the *Sa. islandicus* REY15A. These two ePKs are directly regulated by aCcrs, but the aCcrs binding motif AGTATTAC of ePK2 locates at position +9 downstream of the start codon. Our research reveals that ePK2 exhibits significant cyclic expression (highly expressed in M/G1) in *Sa. islandicus* REY15A [24], consistent with findings in *Sulfolobus acidocaldarius* [7]. Notably, our recent study suggests that cyclically expressed ePK2 influences cell cycle progression[31]. These findings lay the foundation for exploring the role of protein kinases in regulating cell cycle function during early life.

*CRISPR system (G2)*

Although statistical significance was not evident in our functional enrichment analysis, a large number of defense-related genes were indeed enriched among the aCcr2 down-regulated genes as well as among the aCcr2 and aCcr3 up-regulated genes (Figure 4E). Interestingly, CRISPR-associated proteins such as Cbp1, Csa3, Cas3, and Csx1 are downregulated by aCcrs (Table 1 and Table S3). Cren7, a major Sulfolobales chromatin protein, which was found to play an important role in the transcription of CRISPR arrays [34], was also downregulated by aCcrs. EMSA analysis showed that the promoter of Cren7 could be bound by all three aCcrs (Figure S6). Previous analysis of transcriptomes of synchronized *Sa. islandicus* populations also revealed high expression of certain CRISPR-related genes, such as Csa3 and Cbp1, during the G2 phase [21,24]. It is worth noting that Cren7 is also significantly expressed during the S-G2 phase. This evidence demonstrates that aCcrs likely play a role in regulating the

CRISPR defense system.

*Carbohydrate metabolism (S-G2)*

In a previous report, carbohydrate metabolism genes were generally enriched during G2 phase, whereas some carbohydrate transporters were signature genes of the S phase [21]. In eukaryotic cells, cells in G1 and G2 phases undergo an increase in size and energy accumulation in preparation for genome replication and cell division, respectively [11,54]. Carbohydrate metabolism genes were enriched among up-regulated genes in the aCcr2 overexpression cells, implying that aCcr2 likely regulates this process. However, given that aCcr2 functions as a transcriptional repressor, we speculate that this regulation may be indirect. At the same time, carbohydrate metabolism also affects nucleotide and amino acid metabolism, so we speculate that these processes may be influenced by aCcrs in cells and regulated by a variety of complex signals.

### SUPPLEMENTARY FIGURES AND TABLES

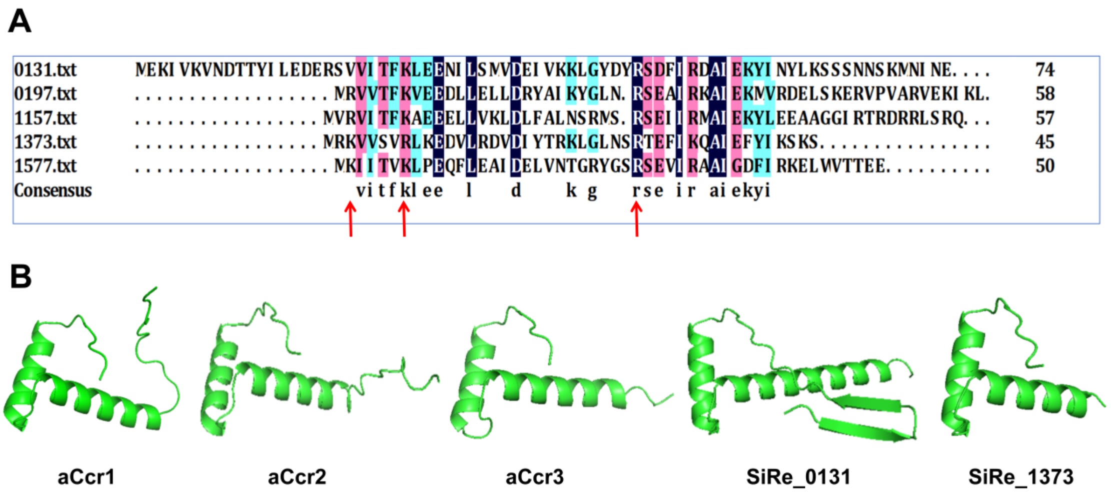


**Figure S1. Multiple sequence alignment and structural prediction of aCcr1 and its homologous proteins using Alphafold2.** (**A**) The protein sequences of aCcr1 (SiRe_0197), aCcr2 (SiRe_1157), aCcr3 (SiRe_1577), SiRe_0131, and SiRe_1373 were aligned using DNAMAN software. Amino acids highlighted by red arrows represent the DNA binding sites previously identified in aCcr1. (**B**) Predicted monomeric protein structures of aCcr1 and its homologs generated by Alphafold3.

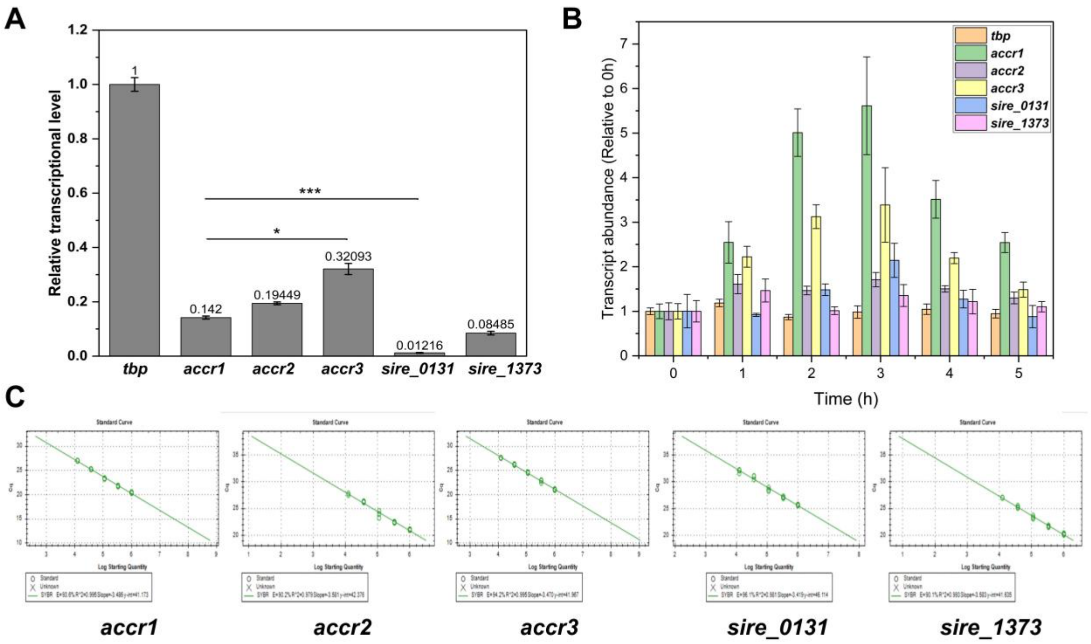


**Figure S2. Expression of *accr1* and its cognate genes.** Overall relative transcription levels (**A**) and expression patterns (**B**) of *accr1* and its cognate genes (*accr2*, *accr3*, *sire_0131*, and *sire_1373*) in the synchronized cells based on the transcriptomic data [24]. *tbp* was used as a reference. Statistical significance was calculated by a One- Sample t-test. *** = p value < 0.0001.** = p value < 0.001. * = p value < 0.01. (**C**) Primer efficiency of qPCR verification for *accr1* and its homologs. Three replicates, five gradients, and a 3-fold dilution gradient were utilized. The primer efficiencies were as follows: 93.6% for accr1, 90.2% for accr2, 94.2% for accr3, 96.1% for sire_0131, and 90.1% for sire_1373, with R2 values greater than or equal to 97.9%.

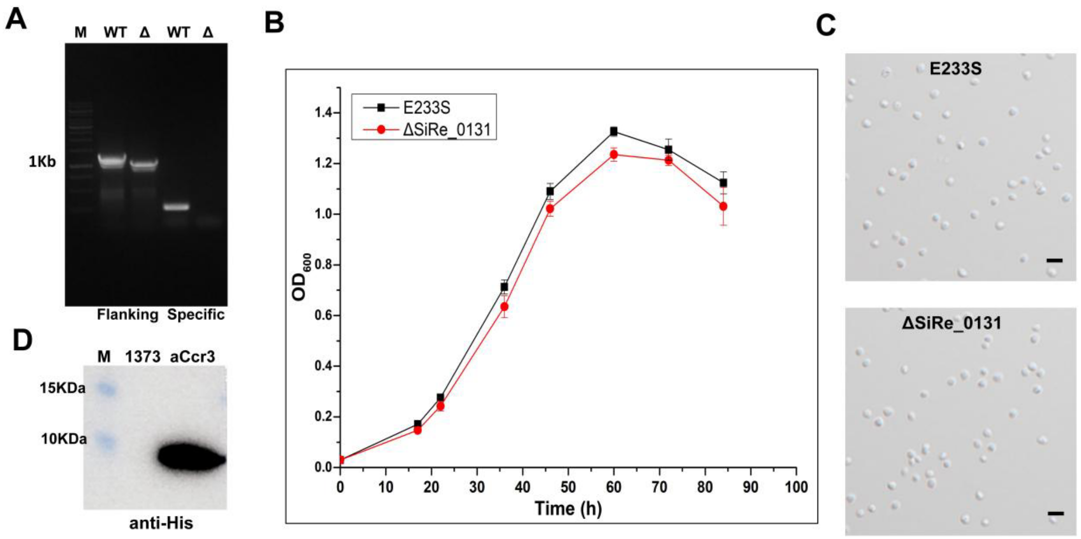


**Figure S3. Phenotypes of Δ*sire_0131* and attempt to overexpress SiRe_1373 in *Sa. Saccharolobus*.** (**A**) PCR validation of *sire_0131* knockout strain using gene flanking primers as well as specific primers. M: marker (**B**) Growth curves of Δ*sire_0131*. E233S was used as a control and the medium was MTSVU. (**C**) Observation of cell morphology of Δ*sire_0131* strain. Scale bars, 2 μm. (**D**) Western blotting to detect whether SiRe_1373 was expressed or not. aCcr3 overexpression strain was used as a positive control.

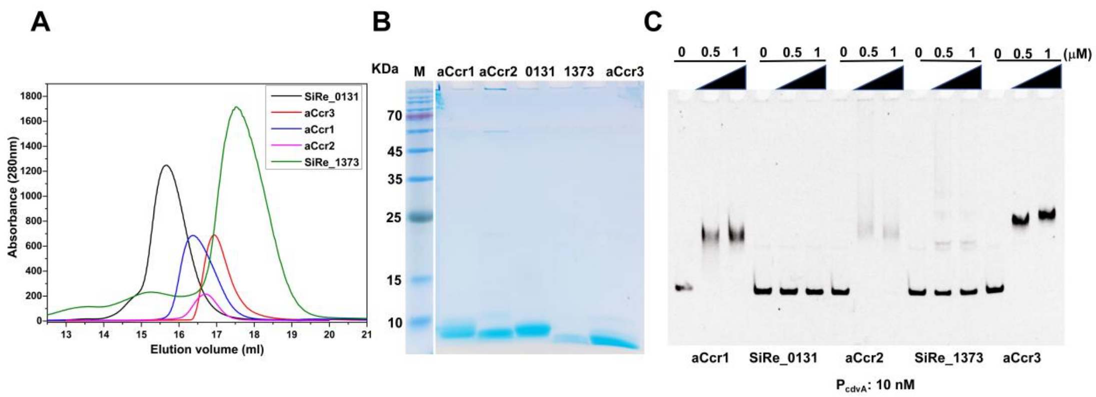


**Figure S4. Purification of aCcr proteins and assay for their binding ability to $P_{cdvA}$.** (**A**) Size exclusion profiles of the purified aCcr1, aCcr2, aCcr3, SiRe_0131, and SiRe_1373. The protein was expressed in *E. coli* and purified by heat treatment, nickel affinity, and gel filtration with a Superdex 200 column as described in the "Materials and Methods". The proteins eluted at 15.6 ml (SiRe_0131, monomeric size 8.76 kDa), 16.4 ml (aCcr1, 6.89 kDa), 16.7 ml (aCcr2, 6.71 kDa), 16.9 ml (aCcr3, 5.77 kDa), and 17.5 ml (SiRe_1373, 5.37 kDa), respetivly, suggesting they form dimers in sollution. (**B**) SDS-PAGE analysis of the recombinant proteins. Purified protein (5 µg) for the aCcrs were loaded for the analysis. The homologs of aCcr1 were expressed and purified as for the wild type aCcr1. M, molecular size marker. (**C**) EMSA for the DNA binding activity of the aCcr1 and homologs. $P_{cdvA}$ was used as the substrate. Each reaction contained 10 nM of the 5'-FAM labelled probe and 0, 0.5 or 1.0 µM proteins.

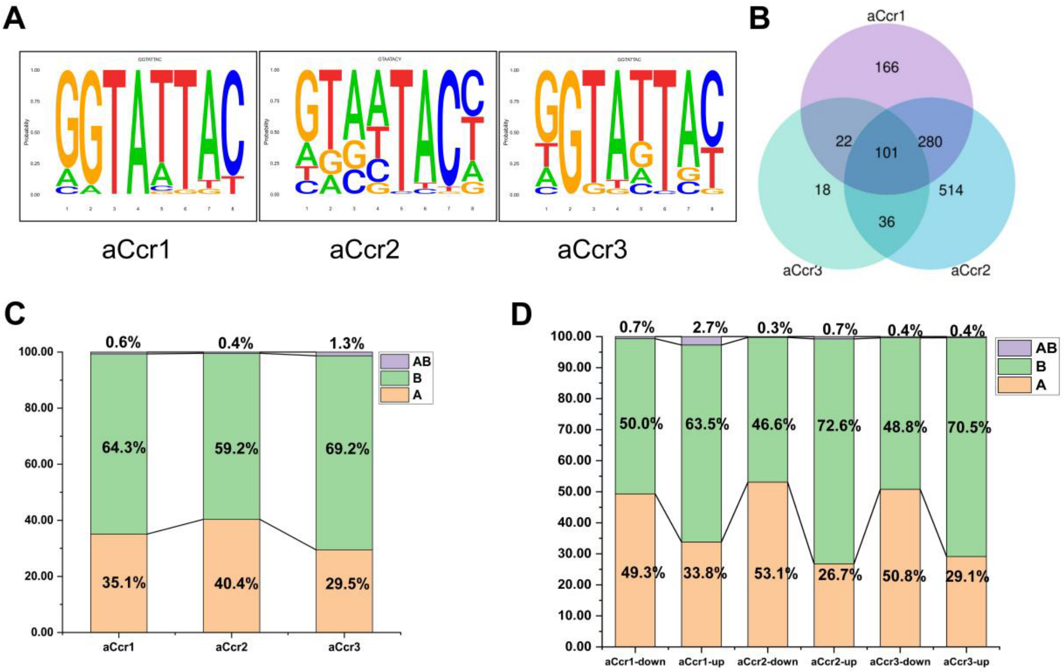


**Figure S5. Statistics of the transcriptomic and ChIP-seq enriched genes. (A)** Sequence logos of the aCcr1, aCcr2 and aCcr3 binding representing MEME predictions of ChIP-seq enriched sequences. (**B**) Venn diagrams of ChIP-seq enriched genes of aCcr1, aCcr2, and aCcr3. (**C**) and (**D**) Chromatin compartment distribution of the ChIP-seq enriched genes (C) and the down-regulated (left) and up-regulated (right) genes by transcriptomic analyses (D). Information on compartmentalization of the genome was extracted from data obtained previously

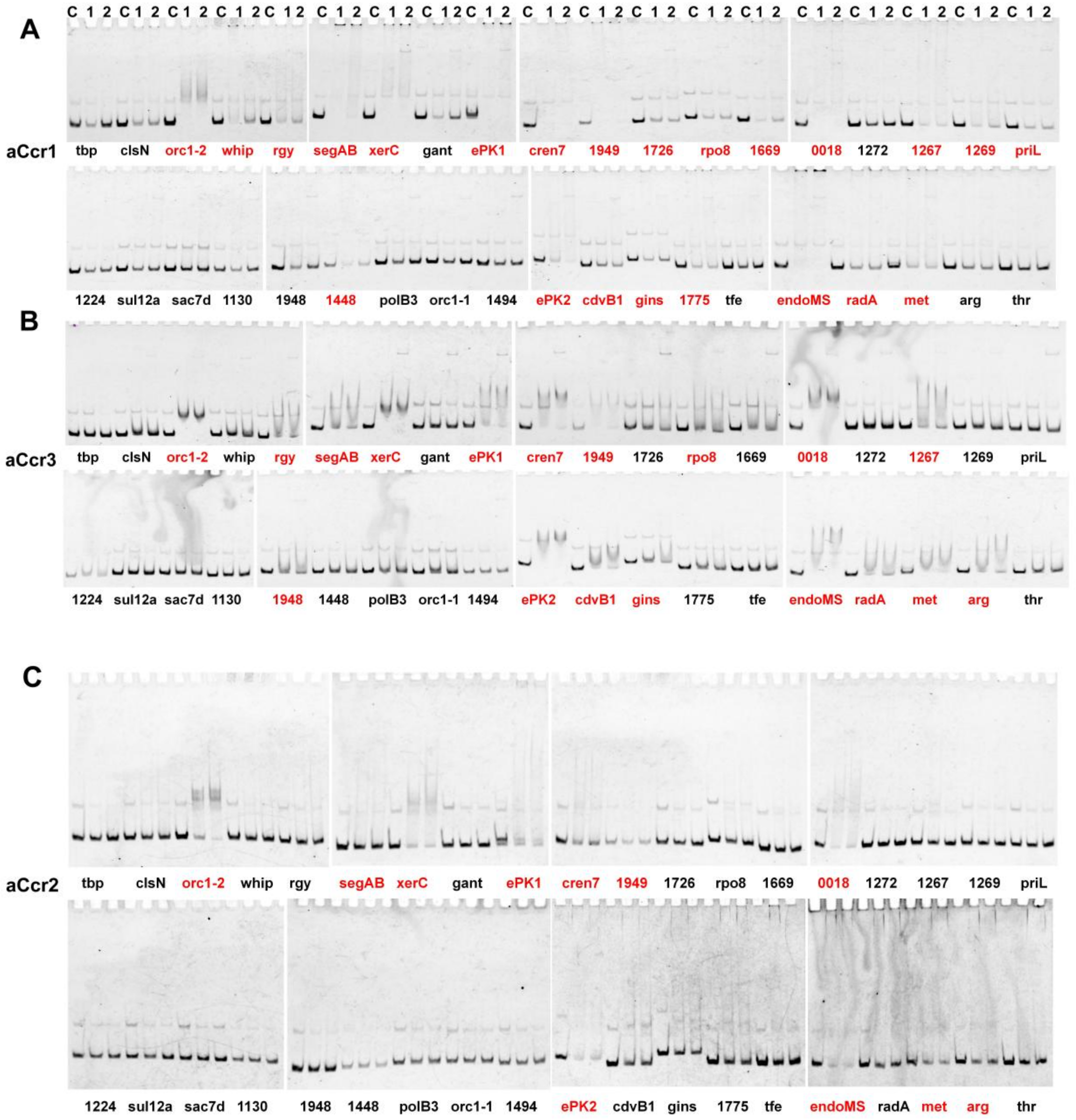


**Figure S6. Promoter binding ability of selected genes based on ChIP-seq and transcriptomic analysis by EMSA.** Thirty-seven promoter fragments were selected from the top down-regulated genes in the aCcr2-overexpress strain. 5’ FAM-DNA fragments containing promoters of interest (100 bp) were used as the substrates. Each reaction contained 10 nM of 5’FAM-labelled probe. C, no protein; 1, 0.5 μM protein; 2, 0.5 μM protein plus 10-fold unlabeled *tbp* promoter DNA. Red labelling indicates those that exhibited direct binding.

**Table 1. Summary of congruently downregulated essential genes in the aCcr overexpression strains based on comparative transcriptomic analysis.**

| Gene identity | Annotation | Transcription Pattern[1] | Promoter binding | Compartment |
|---|---|---|---|---|
| SiRe_0018 | Queuine tRNA-ribosyltransferase related protein | cyclic | yes | A |
| SiRe_0019 | Queuine/archaeosine tRNA-ribosyltransferase | cyclic | yes | A |
| SiRe_0086 | OLD family nuclease | cyclic | yes | A |
| SiRe_1111 | Chromatin protein Cren7 | cyclic | yes | A |
| SiRe_1130 | Uncharacterized protein | N | no | A |
| SiRe_1173 | CdvA | cyclic | yes | A |
| SiRe_1175 | Vps4 | cyclic | no | A |
| SiRe_1226 | Uncharacterized membrane protein | N | no | A |
| SiRe_1448 | Thioredoxin superfamily protein | N | aCcr1 | A |
| SiRe_1949 | AbrB family transcriptional regulator | cyclic | yes | A |

[1] cyclic, cyclic expression. N, no cyclic expression.

**Table S1. Plasmids and strains used in the current study.**

| Strain/Plasmid | Description | References or sources |
|---|---|---|
| *Sulfolobus islandicus* REY15A | Wild type | Contursi et al. |
| *S. islandicus* REY15A (E233S) | *ΔpyrEFΔlacS* | Deng et al. |
| *Escherichia coli* DH5α | Plasmid amplification | Laboratory strain |
| *E. coli* BL21(DE3) codon plus-RIL | Protein expression | Laboratory strain |
| *E. coli* /pET22b-aCcr2-C-His | Expression of WT-aCcr2 having His-tag at the C-terminal | This study |
| *E. coli* /pET22b-aCcr3-C-His | Expression of WT-aCcr3 having His-tag at the C-terminal | This study |
| *E. coli* /pET22b-SiRe_1373-C-His | Expression of WT-SiRe_1373 having His-tag at the C-terminal | This study |
| *E. coli* /pET22b-SiRe_0131-C-His | Expression of WT-SiRe_0131 having His-tag at the C-terminal | This study |
| Sis/pSeSD-aCcr2-C-His | E233S harboring pSeSD-aCcr2-C-His | This study |
| Sis/pSeSD-aCcr3-C-His | E233S harboring pSeSD-aCcr3-C-His | This study |
| Sis/pSeSD-SiRe_1373-C-His | E233S harboring pSeSD-SiRe_1373-C-His | This study |
| Sis/pSeSD-SiRe_0131-C-His | E233S harboring pSeSD-SiRe_0131-C-His | This study |
| pSeSD | A Sulfolobus-E. coli shuttle vector carrying an expression cassette controlled under a synthetic strong promoter ParaS-SD | Peng et al. |
| ΔSiRe_0131 | E233S strain with SiRe_0131 knockout | This study |

### Table S2 Oligonucleotides used as primers in the current study.

| Primer | Sequence [a] (5´-3´) |
|---|---|
| PET22b/pSeSD-aCcr2-*Nde*I-F | GAATGAGGTGAAGCTCATATGTTGGTTAGAGTAATCACTTT |
| **PET22b/pSeSD-aCcr2-*Sal*I-R** | GGCCGCTTGATCAGCGTCGACCTGCCTAGATAGCCGGCGGT |
| PET22b/pSeSD-aCcr3-*Nde*I-F | GAATGAGGTGAAGCTCATATGATGAAAATAATAACGGTA |
| **PET22b/pSeSD-aCcr3-*Sal*I-R** | GGCCGCTTGATCAGCGTCGACCTCTTCTGTAGTTACCCA |
| PET22b/pSeSD-SiRe_0131-*Nde*I-F | GAATGAGGTGAAGCTCATATGGTGGAAAAGATAGTGAAAGT |
| PET22b/pSeSD-SiRe_0131-*Sal*I-R | GGCCGCTTGATCAGCGTCGACTTCATTGATATTCATCTTAC |
| PET22b/pSeSD-SiRe_1373-*Nde*I-F | GAATGAGGTGAAGCTCATATGATGCGAAAAGTAGTAAGTGT |
| PET22b/pSeSD-SiRe_1373-*Sal*I-R | GGCCGCTTGATCAGCGTCGACGCTCTTACTCTTTATATAGA |
| KOaCcr2-Spacer-F | AAGAATATTTTTCTATAGCCATTCTAATTATTTCGCTTCTACT |
| KOaCcr2-Spacer-R | AGCAGTAGAAGCGAAATAATTAGAATGGCTATAGAAAAATATT |
| KOaCcr2-L-F | AAGTACAATTGTGCTGCATGCCAATAAGATAGATGTAGTTA |
| KOaCcr2-R-R | TTAACATATTGGATGCTCGAGATTACTTTAATCTTTCTTAA |
| KOaCcr2-SOE-F | CAATTTATAAAAGAGGAAAACTTACAAGGCCGGCGCTCTA |
| KOaCcr2-SOE-R | TAGAGCGCCGGCCTTGTAAGTTTTCCTCTTTTATAAATTG |
| KOaCcr3-Spacer-F | AAGGAGCAGTTTTTGGAAGCAATAGATGAATTAGTTAATACAG |
| KOaCcr3-Spacer-R | AGCCTGTATTAACTAATTCATCTATTGCTTCCAAAAACTGCTC |
| KOaCcr3-L-F | AAGTACAATTGTGCTGCATGCTACTTTCCAAGAGAAACTAG |
| KOaCcr3-R-R | TTAACATATTGGATGCTCGAGTATCTTCTCTTACATCATAA |
| KOaCcr3-SOE-F | ACTATGCTATCTGAGTTTCTATGATGAAGGCTGGTATTTT |
| KOaCcr3-SOE-R | AAAATACCAGCCTTCATCATAGAAACTCAGATAGCATAGT |

| | |
|---|---|
| KOSiRe_1373-Spacer-F | AAGAAATAGTAGAACTGAGTTCATAAAGCAAGCAATTGAATTC |
| KOSiRe_1373-Spacer-R | AGCGAATTCAATTGCTTGCTTTATGAACTCAGTTCTACTATTT |
| KOSiRe_1373-L-F | AAGTACAATTGTGCTGCATGCTACTAGGTGGAAGCGACTAT |
| KOSiRe_1373-R-R | TTAACATATTGGATGCTCGAGTCTTGACGTCCGGGTATAAT |
| KOSiRe_1373-SOE-F | CTGACAACATTCTATTACATAGGAAACCTTTAACATGCTA |
| KOSiRe_1373-SOE-R | TAGCATGTTAAAGGTTTCCTATGTAATAGAATGTTGTCAG |
| KOSiRe_0131-Spacer-F | AAGCTTATATACTAGAAGATGAACGATCGGTTGTTATAACATT |
| KOSiRe_0131-Spacer-R | AGCAATGTTATAACAACCGATCGTTCATCTTCTAGTATATAAG |
| KOSiRe_0131-L-F | AAGTACAATTGTGCTGCATGCAGAAATAATAATGTAGAAGA |
| KOSiRe_0131-R-R | TTAACATATTGGATGCTCGAGTCCACATCTTATCAACAACC |
| KOSiRe_0131-SOE-F | ATTTATATAGGCGATAATTTAAGTTTAAGTAAACACAAAT |
| KOSiRe_0131-SOE-R | ATTTGTGTTTACTTAAACTTAAATTATCGCCTATATAAAT |
| 0131-Flanking-F | GGAGCTACAATTCCCCTAGGCCC |
| 0131-Flanking-R | CACGTAGCGAGTGAACGCCCAC |
| 0131-Specific-F | GAATGAGGTGAAGCTCATATGGTGGAAAAGATAGTGAAAGT |
| 0131-Specific-R | GGCCGCTTGATCAGCGTCGACTTCATTGATATTCATCTTAC |
| aCcr1-qPCR-F | GGTAGAAGAAGACCTATTAGAG |
| aCcr1-qPCR-R | CCTTAGATAACTCATCCCTTACC |
| aCcr2-qPCR-F | GCAGAGGAGGAACTTTTAGTA |
| aCcr2-qPCR-R | CGGCGGCTTCTTCCAAATATTT |
| aCcr3-qPCR-F | CGGTAAAACTACCAGAGCAGTT |
| aCcr3-qPCR-R | CATAGTTCTTTCCGTATAAAATCACC |

| | |
|---|---|
| 0131-qPCR-F | CTAGAAGATGAACGATCGGTTG |
| 0131-qPCR-R | ATCGTAGCCTAACTTCTTCAC |
| 1373-qPCR-F | GCGAAAAGTAGTAAGTGTAAG |
| 1373-qPCR-R | GGAAGATGTACTGAGAGATGT |
| 16SrRNAqPCR-F | CGCAAGACTGAAACTTAAAGGA |
| 16SrRNAqPCR-R | AGTCAGGCAAGGTCGTTAG |
| TBP-F | GTGGCAACAGTTACGTTAGAG |
| TBP-R | CCTTGGGCTGTTCTAATCTG |
| $P_{cren7}$-F | AGGTCGACTCTAGA<u>GGATCC</u>AACCTTTAATATTACCCTTG |
| $P_{cren7}$-R | ACTTTCTCTATAATACTACT |
| $P_{cdvB1}$-F | AGGTCGACTCTAGA<u>GGATCC</u>CTTTAATGAACGTGTTTGGC |
| $P_{cdvB1}$-R | GGCTAATATATAACTTACCT |
| $P_{orc1\text{-}2}$-F | AGGTCGACTCTAGA<u>GGATCC</u>CTGGGATATCGCTTTTGAGA |
| $P_{orc1\text{-}2}$-R | AACTTTTACCCCAATCTAAA |
| $P_{RadA}$-F | AGGTCGACTCTAGA<u>GGATCC</u>TACCAGTTAAAGATATAGGA |
| $P_{RadA}$-R | CTATTATCACCAGACACTTA |
| $P_{gins\text{-}mcm}$-F | AGGTCGACTCTAGA<u>GGATCC</u>TCATTTATACATTTCTGACC |
| $P_{gins\text{-}mcm}$-R | GCTCTAAGTTTTACTTCAATCACAATC |
| $P_{priL}$-F | AGGTCGACTCTAGA<u>GGATCC</u>ATTTACATATGGCCATGCAG |
| $P_{priL}$-R | TTCCTTACTTATAGTATGTT |
| $P_{1272\text{-}1276}$-F | AGGTCGACTCTAGA<u>GGATCC</u>AAATAGGATACGCCTCAACC |
| $P_{1272\text{-}1276}$-R | AAAAAAAATCCACCTTATTA |

| | |
|---|---|
| **$P_{1267-70}$-F** | AGGTCGACTCTAGAGGATCCACCTACATTGGTCTTTCCTA |
| **$P_{1267-70}$-R** | CTCTTCTCAAAATACTATGC |
| $P_{gyr}$-F | AGGTCGACTCTAGAGGATCCTCAAATAGTCATTCATATAA |
| $P_{gyr}$-R | AGCTAGAAAAAACTTTATGA |
| $P_{orc1-1}$-F | AGGTCGACTCTAGAGGATCCTGACACAACTTACATTTTCC |
| $P_{orc1-1}$-R | CTTTTCTATTGTTTGTGCTA |
| $P_{1129-30}$-F | AGGTCGACTCTAGAGGATCCTCAGAGAACTAGTTTATATT |
| $P_{1129-30}$-R | AACTATTATGTTAGGAAAAAGC |
| $P_{1224-26}$-F | AGGTCGACTCTAGAGGATCCAGTTCATCTAATATGTTCTT |
| $P_{1224-26}$-R | AGCTTATAACTCACTAAATAAAC |
| $P_{1448}$-F | AGGTCGACTCTAGAGGATCCTGTTTTCGCTAACGGTTCTA |
| $P_{1448}$-R | CCCGTATCACTAGTTAAAAGGG |
| $P_{1949}$-F | AGGTCGACTCTAGAGGATCCCATTACTACATCATTCTCAC |
| $P_{1949}$-R | TCTATTCACCATAATACTCTATTAC |
| $P_{clsN}$-F | AGGTCGACTCTAGAGGATCCGTTATTGAGAAAGAGTATTT |
| $P_{clsN}$-R | CTAGTGGTATTAAAGATAAAC |
| $P_{whip}$-F | AGGTCGACTCTAGAGGATCCTAAAACTTGCCAATGAGTAG |
| $P_{whip}$-R | CTTATTATATTAGGCATTATAAC |
| $P_{1494}$-F | AGGTCGACTCTAGAGGATCCATTAATGCATTATATATTTA |
| $P_{1494}$-R | AAAAATGTATTAGTAAAAGAAGG |
| $P_{xerC}$-F | AGGTCGACTCTAGAGGATCCACAATAGTGAAATCAGAAGT |
| $P_{xerC}$-R | GTATGATATATAGTGGAGTTAAG |

| | |
|---|---|
| $P_{1669}$-F | AGGTCGACTCTAGAGGATCCATATTACATAATATTAAATA |
| $P_{1669}$-R | CGCATGTGTATATAAAAGATGATC |
| $P_{rpo8}$-F | AGGTCGACTCTAGAGGATCCAGTCTTATCTATTAGATTAT |
| $P_{rpo8}$-R | GATTCCATCATATTACTATTTAAGTT |
| $P_{1726}$-F | AGGTCGACTCTAGAGGATCCCACAGTATTCTTCACTATTA |
| $P_{1726}$-R | TTTAACTCGTATATATACTTG |
| $P_{TFE}$-F | AGGTCGACTCTAGAGGATCCTTGAATTATTAAGGGAGAGG |
| $P_{TFE}$-R | TCTTTTATCTCTAGCTGGTCTATGG |
| $P_{1775}$-F | AGGTCGACTCTAGAGGATCCCCTTAATTGGCTGAAGACTC |
| $P_{1775}$-R | CTATTTGTTACCTCTTAATGG |
| $P_{GANT}$-F | AGGTCGACTCTAGAGGATCCCTATCAACTAGATCCTTACT |
| $P_{GANT}$-R | ATCAATATCTTTATATAATGTAG |
| $P_{polb3}$-F | AGGTCGACTCTAGAGGATCCGCACGCGTTTGTGATGATAA |
| $P_{polb3}$-R | CATATATTACTAACCAAATGACTGGC |
| $P_{1948}$-F | AGGTCGACTCTAGAGGATCCATAATACTTGCATTAGAGGT |
| $P_{1948}$-R | TTATCTCACTAATAAAGAATC |
| $P_{segAB}$-F | AGGTCGACTCTAGAGGATCCAAAAAACTAGCCTATAAATA |
| $P_{segAB}$-R | GTCTAGACTCTTCTATCTATAACG |
| $P_{sul12a}$-F | AGGTCGACTCTAGAGGATCCGTAAATTTATTACATCATTC |
| $P_{sul12a}$-R | ATATAATAGCTAATCATAGAATCTC |
| $P_{sac7d}$-F | AGGTCGACTCTAGAGGATCCTAATAACTTTAGTTTGCGCT |
| $P_{sac7d}$-R | GTTTCTTCAAGTTATTCTTGAAGG |

| | |
|---|---|
| $P_{\mathit{0018\text{-}19}}$-F | AGGTCGACTCTAGAGGATCCGTGATATTTTCGTTAACTAA |
| $P_{\mathit{0018\text{-}19}}$-R | TTTTCCTGGTATTGGTATTAC |
| $P_{\mathit{ePK1}}$-F | AGGTCGACTCTAGAGGATCCAAGTAAGAAAATTAGAGTTA |
| $P_{\mathit{ePK1}}$-R | GGTATTACTATTTTACTGGTAAC |
| $P_{\mathit{ePK2}}$-F | AGGTCGACTCTAGAGGATCCAAGTGTGGTGTCCATAATCT |
| $P_{\mathit{ePK2}}$-R | CTGTAATACTAACTGCACATTAAC |
| $P_{\mathit{tbp}}$-F | AGGTCGACTCTAGAGGATCCATACTTATTTAAAAGAGGGA |
| $P_{\mathit{tbp}}$-R | ATTTACAATGGGTTTATACGATACC |
| $P_{\mathit{nucS}}$-F | AGGTCGACTCTAGAGGATCCTATTAGCCCCAAATACAAGG |
| $P_{\mathit{nucS}}$-R | CACAGTATTACTTATCATCAATGC |
| $P_{\mathit{Met}}$-F | AGGTCGACTCTAGAGGATCCAATAGCTCAAGAATTAAGTT |
| $P_{\mathit{Met}}$-R | CCGCTATTCTACCCCGCTAT |
| $P_{\mathit{Arg}}$-F | AGGTCGACTCTAGAGGATCCTTTTGAAGTCTAGTTAAAAT |
| $P_{\mathit{Arg}}$-R | GGTCCCCTTATGATACTCTC |
| $P_{\mathit{Thr}}$-F | AGGTCGACTCTAGAGGATCCGCAATAATTAAAGGTTCAGA |
| $P_{\mathit{Thr}}$-R | GCGGCATCATTTTACTATAT |

[a] The underlined denote sites of restriction enzymes.

**Table S3. Summary of congruently downregulated non-essential genes in the aCcr overexpression strains based on comparative transcriptomic analysis.**

| Gene identity | Annotation | Transcription Pattern[1] | Promoter binding[2] | Compartment |
|---|---|---|---|---|
| SiRe_1547 | CRISPR repeat-binding protein(cbp1) | cyclic | no | A |
| SiRe_0764 | CRISPR-associated, Csa3 | cyclic | aCcr1,2 | A |
| SiRe_0421 | CRISPR-associated helicase Cas3 | N | aCcr1 | B |
| SiRe_2599 | CARF domain containing protein, contains HTH a nd HEPN domains，Csx1 | N | no | A |
| SiRe_1740 | Cdc6-related protein, Orc1-1 | cyclic | no | A |
| SiRe_1861 | PolB1-binding proteins PBP1, C2H2-type zinc finger | N | no | A |
| SiRe_0087 | Predicted membrane protein, DUF131 family | cyclic | yes | A |
| SiRe_2057 | Uncharacterized membrane protein | cyclic | yes | B |
| SiRe_0025 | NucS/EndoMS | N | yes | A |
| SiRe_0017 | HerA helicase | cyclic | yes | A |
| SiRe_1224 | RecA/RadA recombinase | N | no | A |
| SiRe_1605 | Lhr-like helicase with C-terminal Zn finger domain | cyclic | yes | A |
| SiRe_0691 | Thermopsin-like protease | N | yes | B |
| SiRe_0931 | Thermopsin-like protease | N | aCcr2,3 | B |
| SiRe_2056 | Membrane associated serine/threonine protein kina se | cyclic | yes | B |

| | | | | |
|---|---|---|---|---|
| SiRe_2030 | serine/threonine protein kinase | cyclic | yes | B |
| SiRe_0623 | Uncharacterized membrane protein | cyclic | yes | B |
| SiRe_0624 | Sulfocyanin | cyclic | yes | B |
| SiRe_0265 | ATPase involved in chromosome partitioning, ParA family | N | aCcr2 | B |
| SiRe_1963 | segC | N | aCcr2 | AB |
| SiRe_1239 | Zn-dependent alcohol dehydrogenase | N | aCcr2 | A |
| SIRE_RS02790 | IS1 family transposase | N | aCcr2 | B |
| SiRe_1973 | Glycine/D-amino acid oxidase (deaminating) | N | no | B |
| SiRe_1238 | Asp-tRNAAsn/Glu-tRNAGln amidotransferase A subunit or related a midase | N | aCcr2 | A |
| SiRe_0420 | Ser-tRNA(Ala) deacylase AlaX (editing enzyme) | N | aCcr1 | B |
| SiRe_0688 | Amino acid transporter | cyclic | aCcr1 | B |
| SiRe_2216 | L-alanine-DL-glutamate epimerase or related enzyme of enolase superfamily | cyclic | yes | B |
| SiRe_1464 | DUF4352 domain-containing protein | N | yes | A |
| SiRe_2137 | DUF998 domain-containing protein | cyclic | yes(intragenic) | B |
| SiRe_1499 | 5,10-methylenetetrahydrofolate reductase | N | aCcr2 | A |
| SiRe_0917 | HEPN domain containing protein | N | no | B |
| SiRe_1104 | Uncharacterized helical repeat protein, DUF1955 f amily | N | no | A |

| | | | | |
|---|---|---|---|---|
| SiRe_1086 | ABC-type sugar transport system, permease component | N | aCcr2,3 (intragenic) | A |
| SiRe_1087 | ABC-type sugar transport system, permease component | N | aCcr1 (intragenic) | A |
| SiRe_1088 | ABC-type trehalose transport system, periplasmic component | N | aCcr2,3 | A |
| SiRe_2281 | ABC-type transport system, periplasmic component | N | yes | B |
| SiRe_1004 | Uncharacterized protein | N | no | B |
| SiRe_1624 | Sodium/hydrogen exchanger | cyclic | yes | A |
| SiRe_2009 | Threonine dehydrogenase or related Zn-dependent dehydrogenase | cyclic | yes | B |
| SiRe_2201 | Gluconolactonase | cyclic | Yes (intragenic) | B |

[1] cyclic, cyclic expression. N, no cyclic expression.

[2] yes, there are peaks in the promoter in both aCcr1,2,3 ChIP-seq data. No, there are no peaks in all the aCcr1,2,3 ChIP-seq data. intragenic, Peak is located inside the gene.

**Table S4**. **Summary of congruently upregulated genes in the aCcr1, aCcr2, and aCcr3 overexpression strains based on comparative transcriptomic analysis.**

| Gene identity | Annotation | Transcription Pattern[1] | Promoter binding | Compartment |
|---|---|---|---|---|
| SiRe_0906 | RHH/CopG family DNA binding protein | cyclic | yes (intragenic) | B |
| SiRe_1524 | Radical SAM superfamily enzyme | N | yes (intragenic) | A |
| SiRe_0924 | HEPN domain containing protein | N | no | B |
| SiRe_0291 | Zn-dependent alcohol dehydrogenase | N | no | B |
| SiRe_0292 | Acetyl-CoA acetyltransferase | cyclic | yes (intragenic) | B |
| SiRe_0495 | Uncharacterized membrane protein, DUF973 family | cyclic | no | B |
| SiRe_2324 | Xanthine and CO dehydrogenase maturation factor, XdhC/CoxF family | cyclic | no | B |
| SiRe_0133 | ATPase involved in chromosome partitioning, ParA family | N | yes (intragenic) | A |
| SiRe_2636 | Predicted Fe2+/Mn2+ transporter, VIT1/CCC1 family | N | aCcr1 (intragenic) | B |
| SiRe_0742 | Isocitrate lyase | N | no | B |
| SiRe_0290 | Alpha/beta superfamily hydrolase | cyclic | no | B |
| SiRe_2533 | N-acetylglucosamine-6-phosphate deacetylase | N | aCcr1 (intragenic) | B |
| SiRe_0481 | Dienelactone hydrolase or related enzyme | N | no | B |
| SiRe_t0001 | tRNA-Leu | N | aCcr2,3 | A |
| SiRe_0132 | ATPase involved in chromosome partitioning, ParA family | N | no | A |
| SiRe_2541 | Uncharacterized membrane protein | N | no | B |
| SiRe_2567 | Aerobic-type carbon monoxide dehydrogenase, small subunit CoxS/CutS homolog | cyclic | no | A |
| SiRe_0807 | Acetylornithine deacetylase/Succinyl-diaminopimelate desuccinylase or related deacylase | N | yes | B |
| SiRe_2540 | Uncharacterized protein | N | no | B |

| | | | | |
|---|---|---|---|---|
| SiRe_2568 | Aerobic-type carbon monoxide dehydrogenase, middle subunit CoxM/CutM homolog | cyclic | no | A |
| SiRe_2306 | Uncharacterized membrane protein | N | no | B |

[1] cyclic: cyclic expression. N: no cyclic expression. [2] yes: there are peaks in the promoter in both aCcr1,2,3 ChIP-seq data. No: there are no peaks in all the aCcr1,2,3 ChIP-seq data. intragenic: Peak is located inside the gene.

**Supplementary Table S5** Functional enrichment analysis of genes upregulated or downregulated by aCcr1, aCcr2 and aCcr3 and their ChIP-seq data.

**Supplementary Table S6** Comparative transctiptomic data of the strains overexpressing aCcr1, aCcr2 and aCcr3. The strain carrying the empty vectord was used as the control.

**Supplementary Table S7** ChIP-seq enrichement data of aCcr2, the narrow peaks by ChIPseeker_annotation.

**Supplementary Table S8** ChIP-seq enrichement data of aCcr3, the narrow peaks by ChIPseeker_annotation.
